# Micronuclei arising due to loss of KIF18A form stable micronuclear envelopes and do not promote tumorigenesis

**DOI:** 10.1101/2020.11.23.394924

**Authors:** Leslie A. Sepaniac, Whitney Martin, Louise A. Dionne, Timothy M. Stearns, Laura G. Reinholdt, Jason Stumpff

## Abstract

Micronuclei, whole or fragmented chromosomes which are spatially separated from the main nucleus, are strongly associated with genomic instability and have been identified as drivers of tumorigenesis. Paradoxically, *Kif18a* mutant mice produce micronuclei due to unaligned chromosomes *in vivo* but do not develop spontaneous tumors, raising questions about whether all micronuclei contribute similarly to genomic instability and cancer. We report here that micronuclei in *Kif18a* mutant mice form stable nuclear envelopes. Challenging *Kif18a* mutant mice via deletion of the *Trp53* gene led to formation of thymic lymphoma with elevated levels of micronuclei. However, loss of *Kif18a* had modest or no effect on survival of *Trp53* homozygotes and heterozygotes, respectively. To further explore micronuclear envelope stability in *KIF18A* KO cells, we compared micronuclei induced via different insults in cultured cells. Micronuclei in *KIF18A* KO cells form stable nuclear envelopes characterized by increased recruitment of core and non-core nuclear envelope components and successful expansion of decondensing chromatin compared to those induced by microtubule drug washout or exposure to radiation. We also observed that lagging chromosomes, which lead to micronucleus formation, were positioned closer to the main chromatin masses, and further from the central spindle, in *KIF18A* KO cells. Our studies provide *in vivo* support to models suggesting that micronuclear fate depends on the sub-cellular location of late lagging chromosomes and suggest that not all micronuclei actively promote tumorigenesis.

## Introduction

Micronuclei contain chromosomes which are excluded from the main nucleus and are used clinically as a biomarker to evaluate genomic instability (Fenech and Morley, 1985; Tolbert et al., 1992; Dertinger et al., 1996; Fenech, 2000; Bonassi et al., 2007; Imle et al., 2009; Fenech et al., 2011; Luzhna et al., 2013). Micronuclei are widely associated with chromosomally unstable tumors and poor patient prognosis (Bonassi et al., 2007; Imle et al., 2009; Fenech et al., 2011; Luzhna et al., 2013). Growing evidence demonstrates that micronuclei are not only passive markers, but also active drivers of genomic instability – though the specific conditions required for this transformation are not fully elucidated (Stephens et al, 2011; Rausch et al., 2012; Holland and Cleveland, 2012; Crasta et al., 2012; Nones et al., 2014; Zhang et al., 2015; Luijten et al., 2018).

Micronuclei can arise due to various errors occurring during the cell cycle, including improper attachments between microtubules and kinetochores, DNA replication errors, and unrepaired DNA damage (Fenech and Morley, 1985; Cimini et al., 2001; Hoffelder et al., 2004; Crasta et al., 2012). Chromosomes which become micronucleated can be whole or fragmented, and DNA content can be centromere-containing or acentric, with different mechanisms of micronucleus formation leading to varying levels of damage to the micronucleated DNA content (Ding et al., 2003; Hoffelder et al., 2004; Terradas et al., 2009; Terradas et al., 2010; Huang et al., 2011, Crasta et al., 2012, Hatch et al., 2013, Zhang et al., 2015, Liu et al., 2018).

There are two widely accepted, non-mutually exclusive mechanisms that explain how a micronucleated chromosome may lead to genomic instability. First, cells entering cell division with a micronucleus can become fragmented and result in severe, localized DNA rearrangements to the micronucleated chromosome (Crasta et al., 2012; Holland and Cleveland, 2012; Jones and Jallepalli, 2012; Zhang et al., 2015). This catastrophic process, termed chromothripsis, has been identified as an early event in tumorigenesis, and has been elegantly demonstrated in experiments pairing long-term imaging with single-cell whole genome sequencing in cultured cells (Zhang et al., 2015). Further, loss of micronuclear envelope integrity can also lead to genomic instability by exposing the chromosome to damage in the cell’s cytoplasm (Hoffelder et al., 2004; Hatch et al., 2013; Zhang et al., 2015, Shah et al., 2017). While these separate and distinct cellular events may follow one another along a shared pathway, this does not occur in all cases (Hatch et al., 2013, Zhang et al., 2015; He et al., 2019).

More recently, micronuclei have been demonstrated to form as a result of chromosome alignment defects during mitosis (Fonseca et al., 2019). In cultured human cells lacking the function of the kinesin KIF18A, chromosomes fail to properly align at the mitotic spindle equator, segregate in a disordered fashion, and display an increased probability of forming micronuclei (Fonseca et al., 2019). Furthermore, mice with inactivating mutations in *Kif18a* form micronuclei *in vivo,* with micronuclear incidences significantly elevated from levels of spontaneously occurring micronuclei in wild type mice. Paradoxically, *Kif18a* mutant mice do not develop tumors spontaneously and have been shown to be resistant to induced colitis-associated colorectal cancer (Zhu et al., 2013). These results raise a number of questions about the conditions under which micronuclei might induce genomic instability and tumorigenesis *in vivo. Kif18a* mutant mice are a useful system for studying the effects of micronuclei *in vivo* since the level of aneuploidy observed in these mice is low, with no apparent increase in aneuploidy detected in mouse embryonic fibroblasts, allowing separation of effects due to micronucleation from those caused by widespread aneuploidy (Czechanski et al., 2015; Fonseca et al., 2019).

It is possible that micronuclei in *Kif18a* mutant mice minimally impact genomic stability due to p53-dependent cell cycle arrest or maintenance of micronuclear envelope stability. Activation of p53 in micronucleated cells has been observed to cause cell-cycle arrest in the subsequent G1 (Uetake and Sluder, 2010; Santaguida, et al., 2017, Thompson and Compton, 2010), and micronucleated KIF18A KO cells are subject to a p53-dependent cell cycle arrest in culture (Fonseca et al., 2019). It is plausible, then, that micronuclei produced due to loss of *Kif18a* do not contribute to tumorigenesis in mice because a p53-dependent pathway prevents micronucleated cells from dividing further. This possibility is consistent with the results of *in vitro* studies that demonstrated the contribution of micronuclei to genomic instability over repeated divisions where experiments were carried out in cell lines lacking p53 activity (Crasta et al., 2012, Hatch et al., 2013, Zhang et al., 2015, Liu et al., 2018, Soto et al., 2018). In addition, recent studies have demonstrated that not all micronuclei undergo nuclear envelope rupture (Liu et al., 2018; He et al., 2019). Thus, it is also possible that micronuclei in *Kif18a* loss of function cells form stable nuclear envelopes, which could reduce their negative impact on genomic stability.

To investigate the effects of *p53* status and micronuclear envelope stability on the impact of micronuclei *in vivo*, we developed a mouse model lacking *Kif18a* and p53 function. We found that micronuclei arising due to chromosome unalignment formed robust micronuclear envelopes which rupture less frequently than those surrounding micronuclei formed due to improper kinetochore-microtubule attachments. These data indicate that the type of insult which leads to micronucleus formation can impact resulting micronuclear stability, and we speculate that these differences influence the severity of overall risk to genome integrity.

## Results

### Loss of *Kif18a* increases micronuclei in both normal tissues and thymic lymphomas of p53-null mice

Mice homozygous for the *Kif18a* mutation *gcd2* (germ cell depletion 2) lack KIF18A activity and form micronuclei due to chromosome alignment defects *in vivo* (Czechanski et al., 2015; Fonseca et al., 2019). While *Kif18a^gcd2/gcd2^* mice are infertile due to mitotic defects during embryonic germline development, they do not develop spontaneous tumors (Czechanski et al., 2015).

Analyses of KIF18A KO RPE1 cells indicated that micronucleated cells rarely entered mitosis (Fonseca et al., 2019). This arrest was at least partially dependent on p53, consistent with other reports of cell cycle arrest following micronucleation. Thus, we reasoned that a p53-dependent mechanism could limit the impact of micronuclei on tumor induction or development in *Kif18a* mutant mice (Sablina 1998, Thompson and Compton, 2010; Fonseca et al., 2019). To investigate this possibility, we crossed *Kif18a^gcd2/+^* mice with mice heterozygous for a p53 null mutation (*Trp53^tm1^ ^Tyj/+^*) (Fig. 1A).

**Figure 1:**
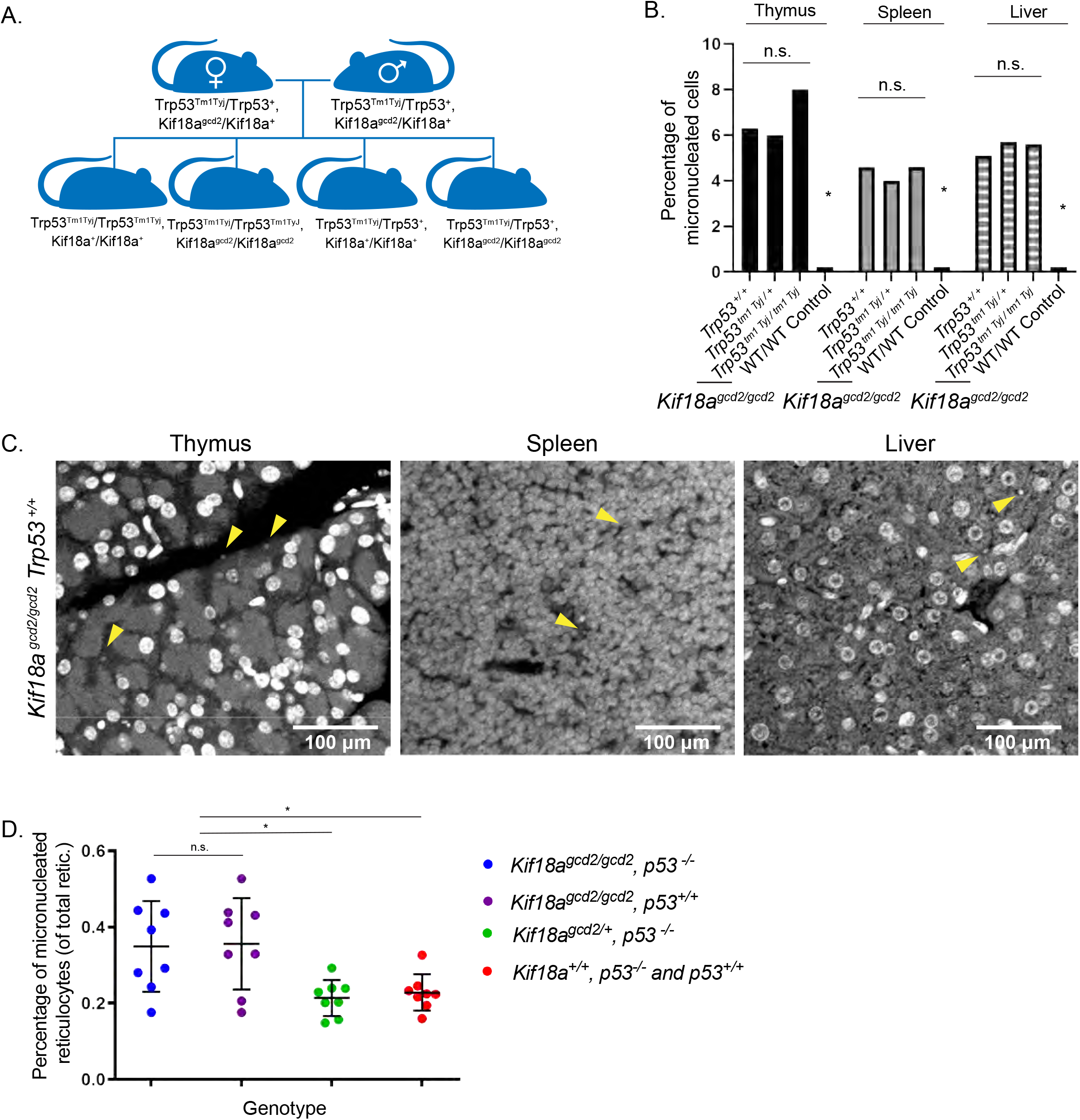
*Kif18a* mutant mice display similarly elevated levels of micronuclei in healthy tissues, regardless of p53 status. **(A)** Schematic of cross used to generate *Kif18a^gcd2^*, *Trp53^tm1^ ^Tyj^* mice. **(B)** Quantification of micronucleated cells, as observed by Hoechst stain, observed in thymus, spleen, and liver tissues from healthy individuals homozygous for the *Kiif18A^gcd2^* mutation, and with wildtype *Trp53, Trp53^tm1^ ^Tyj/+^,* or *Trp53^tm1^ ^Tyj/tm1^ ^Tyj^.* n=3 tissue types from one biological sample per each genotype was scored. Percentages are the average from 2 independent counts of each tissue. Micronucleated cell counts were as follows: *Kif18a^gcd2/gcd2^, Trp53^+/+^* 83/1317 in thymus, 119/2587 in spleen, 57/1115 in liver; *Kif18a^gcd2/gcd2^, Trp53^+/tm1^ ^Tyj^* 120/1496 thymus, 68/1468 spleen, 41/735 liver; *Kif18a^gcd2/gcd2^, Trp53^tm1^ ^Tyj/tm1^ ^Tyj^* 36/602 thymus, 97/2410 spleen, 46/811 liver. (Table S1.) Thymus, p = 0.11, n.s.; spleen p = 0.54, n.s.; liver p = 0.84, n.s. Indicated p values were calculated by χ2 analysis. **(C)** Representative images of micronuclei (indicated by yellow arrowhead) observed in healthy (left to right) thymus, spleen, and liver tissue sections from a *Kif18a^gcd2/gcd2^*, p53^+/+^ mouse. **(D)** Plot showing percentages of micronucleated reticulocytes (of total reticulocytes) quantified via peripheral blood assay from male and female mice of genotypes: *Kif18a^gcd2/gcd2^, Trp53^tm1^ ^Tyj/tm1^* ^Tyj^ (n=8); *Kif18a^gcd2/gcd2^, Trp53^+/+^* (n=8); *Kif18a^gcd2/+^, Trp53^tm1^ ^Tyj/tm1^* ^Tyj^ (n=8); *Kif18a^+/+^, Trp53^tm1^ ^Tyj/tm1^* ^Tyj^ and *Trp53^+/+^* (n=8). Data points indicate individual biological replicates. Error bars represent standard deviation. Statistical analysis was performed using pairwise ANOVA comparisons of means, alpha 0.05 (* p < 0.01).

Micronuclei in *Kif18a* mutant mice were previously quantified in red blood cells via flow cytometry (Fonseca et al., 2019). To confirm that micronuclei are present in other tissues, we analyzed thymus, spleen, and liver tissues from mice carrying *Kif18a* mutations in the presence or absence of *Trp53*. As expected, *Kif18a^gcd2/gcd2^* mice displayed elevated levels of micronuclei in all tissues compared to littermate controls (Fig. 1B-C and Table S1). Healthy thymus, spleen, and liver tissues from *Kif18a^gcd2/gcd2^*, *Trp53 ^tm1^ ^Tyj/tm1 Tyj^* mice exhibited similar percentages of micronucleated cells as *Kif18a^gcd2/gcd2^* mice (Fig. 1C). Additionally, analyses of micronucleated reticulocytes via flow cytometry also indicated that the percentage of micronucleated cells was not affected by p53 *in vivo* (Fig. 1D).

Mice homozygous or heterozygous for null mutations in *Trp53* develop a spectrum of tumors, with a predominance of thymic lymphoma (Jacks et al., 1994). Consistent with this, *Kif18a^gcd2/gcd2^, Trp53^tm1^ ^Tyj/tm1^ ^Tyj^* homozygous mutant mice developed tumors within 3 months, with the vast majority exhibiting thymic lymphoma (78%). To investigate whether the prevalence of micronuclei found within tumor tissues varied among *Kif18a* mutant and *Trp53* mutant animals, we analyzed primary thymic lymphoma sections stained with Hoechst to label DNA. We observed that tumors from *Kif18a^gcd2/gcd2^ , Trp53^tm1^ ^Tyj/tm1^ ^Tyj^* mice exhibited elevated levels of micronucleated cells compared to those from *Kif18a^+/+^ , Trp53^tm1^ ^Tyj/tm1^ ^Tyj^* mice (p < 0.001; Fig. 2A-B).

**Figure 2:**
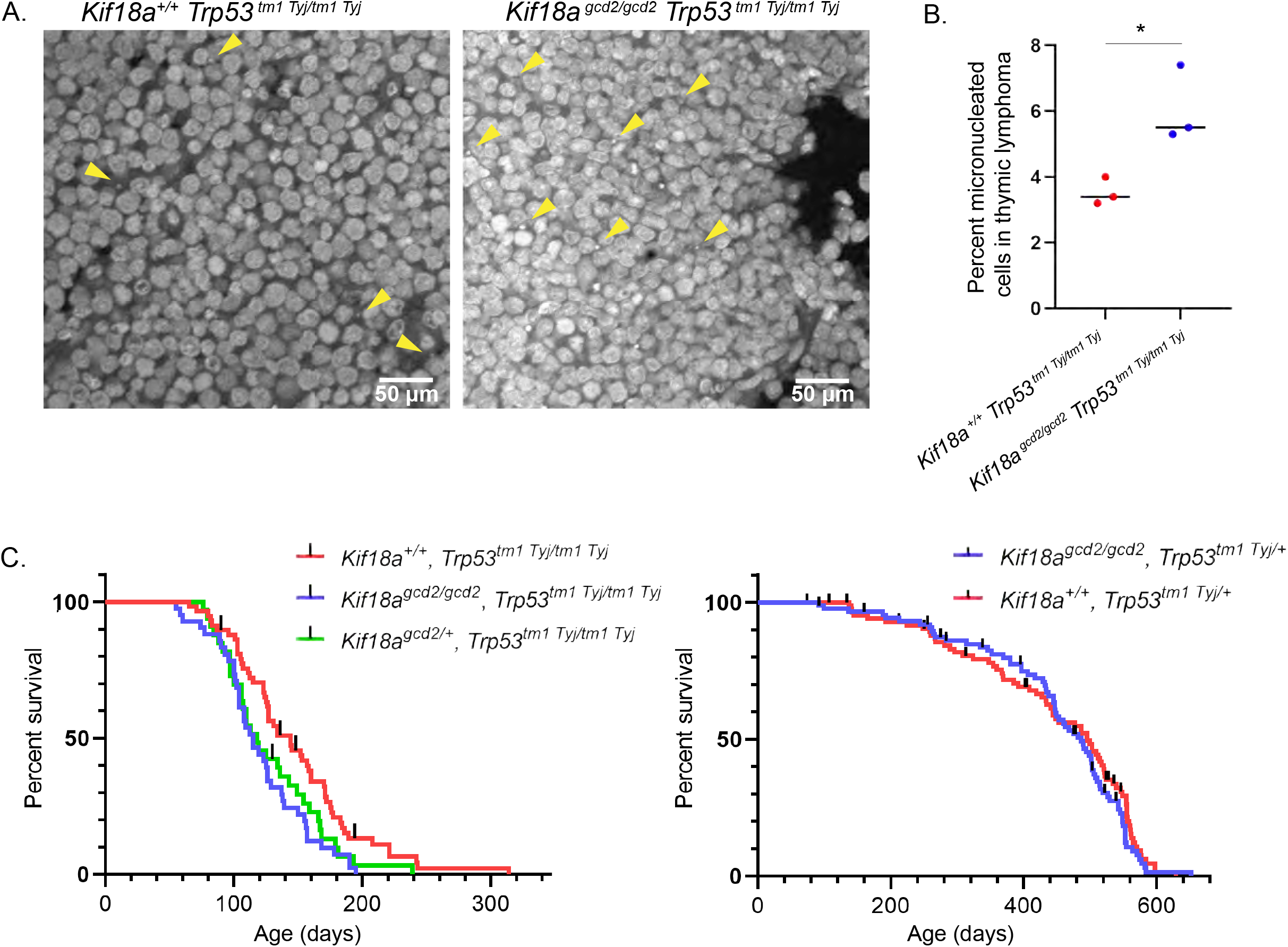
Loss of *Kif18a* function increases the percentage of micronucleated cells in tumors caused by *Trp53* mutation but only modestly reduces survival. **(A)** Representative images of micronuclei (yellow arrowheads) observed in thymic lymphoma tumor sections, stained via Hoechst, from *Kif18a^gcd2/gcd2^, Trp53^tm1^ ^Tyj/tm1^* ^Tyj^ and *Kif18a^+/+^, Trp53^tm1^ ^Tyj/tm1^* ^Tyj^ mice. **(B)** Plot showing the percentage of micronucleated cells observed in sectioned thymic lymphoma from the indicated genotypes. Data points represent individual biological samples. n=3 biological replicates per genotype; *Kif18a^+/+^, Trp53^tm1^ ^Tyj/tm1^* ^Tyj^ n= 3099 total cells scored; *Kif18a^gcd2/gcd2^, Trp53^tm1^ ^Tyj/tm1^* ^Tyj^ n = 4210 total cells scored for micronuclei; * p < 0.001; (Table S2). Statistical comparison was made via χ2 analysis. **(C)** Kaplan Meier survival curves for the indicated genotypes (Left) Loss of *Kif18a* significantly reduced survival of *Trp53^tm1^ ^Tyj/tm1^ ^Tyj^* mice, with an average reduction in survival of 20 days (*Kif18a^+/+^, Trp53^tm1^ ^Tyj/tm1^* ^Tyj^ n = 58; *Kif18a^gcd2/gcd2^, Trp53^tm1^ ^Tyj/tm1^* ^Tyj^ n = 41; *Kif18a^gcd2/+^, Trp53^tm1^ ^Tyj/tm1^* ^Tyj^ n = 33; * p = 0.01). (Right) Loss of *Kif18a* did not significantly reduce the survival of *Trp53^tm1^* ^Tyj/+^ mice (*Kif18a^gcd2/gcd2^, Trp53^tm1^ ^Tyj/+^* n = 88; *Kif18a^+/+^, Trp53^tm1^ ^Tyj/+^* n = 87; p = 0.4284, n.s.). Black lines represent censored datapoints. Indicated p values were obtained by performing log rank analysis of mean survival time and a Wang Allison test of maximal lifespan.

### Loss of *Kif18a* minimally affects the survival of *Trp53* mutant mice

If the elevated levels of micronuclei observed in *Kif18a^gcd2/gcd2^, Trp53^tm1^ ^Tyj/tm^ ^1^ ^Tyj^* accelerated tumorigenesis, we reasoned that this effect would reduce survival. We did find that mice homozygous for both *Kif18a* and p53 mutations had a small but significant reduction in survival compared to p53 null littermates with wild type *Kif18a* (p = 0.01; Fig. 2C, left). The reduced survival of the double mutants could be explained by (1) an increase in tumor development that occurs as a result of the *Kif18a* mutation, (2) an interaction between the *Kif18a* null genotype and the genetic background differences introduced by the cross, or (3) a slightly reduced ability of *Kif18a* null mice to cope with rapid tumorigenesis. To help distinguish between these possibilities, we tested the effects of *Kif18a* loss of function on survival of p53 heterozygotes, which exhibit slower tumor development. Within the p53 heterozygous population, there was no significant difference in survival between *Kif18a^gcd2/gcd2^* and *Kif18a^+/+^* animals (p = 0.4284; Fig. 2C, right). These results demonstrate that reduced survival in the *Kif18a* null, p53 mice cannot be attributed to genetic background effects on the *Kif18a* null genotype. Furthermore, micronuclei formed due to the absence of *Kif18a* are unlikely to contribute strongly to a reduction in survival, since micronucleated cell frequency is similarly high in both p53 homozygotes and heterozygotes lacking *Kif18a* function (Fig. 1C and Table S1). Therefore, the reduced survival observed in *Kif18a^gcd2/gcd2^ , Trp53^tm1^ ^Tyj/tm1^ ^Tyj^* mice is likely due to a reduced ability of *Kif18a* mutants to cope with rapid tumorigenesis.

### Micronuclear envelopes in normal tissues of *Kif18a^gcd2/gcd2^, Trp53^tm1 Tyj/tm1 Tyj^* mice are stable, but those in tumor cells are disrupted

Micronuclear envelope instability has been reported to contribute significantly to genomic instability (Hatch et al., 2013; Shah et al., 2017; Zhang et al., 2015; Liu et al., 2018). Previous i*n vitro* studies indicate that micronuclear envelopes are often incomplete, lacking the appropriate and expected density, deposition, or diversity of some nuclear envelope components (Hatch et al., 2013; Liu et al., 2018). To analyze micronuclear envelopes in *Kif18a* mutant mice, sections from liver, spleen, and thymus were stained for the core nuclear envelope protein lamin A/C and DNA (Fig. 3A). Micronuclei surrounded by continuous lamin A/C signal were considered to have intact nuclear envelopes, while those containing gaps in or completely lacking lamin A/C were considered to have ruptured nuclear envelopes. We found that the majority of micronuclear envelopes remained intact (only 3-17% rupture, varying by tissue type) and did not exhibit evidence of rupture within healthy tissues from *Kif18a* mutant mice (Fig. 3B and Table S1). In contrast, micronuclei found in tissues from WT mice were rare but usually exhibited evidence of rupture (100% of micronuclei in thymus, n=1, were ruptured; 50% of micronuclei in spleen, n=2, were ruptured; 100% of micronuclei in liver, n=1, were ruptured). The incidence of micronuclear envelope rupture was also uniformly low regardless of *Trp53* allele status. Thus, micronuclei formed following loss of KIF18A function *in vivo* appear to have stable nuclear envelopes.

**Figure 3:**
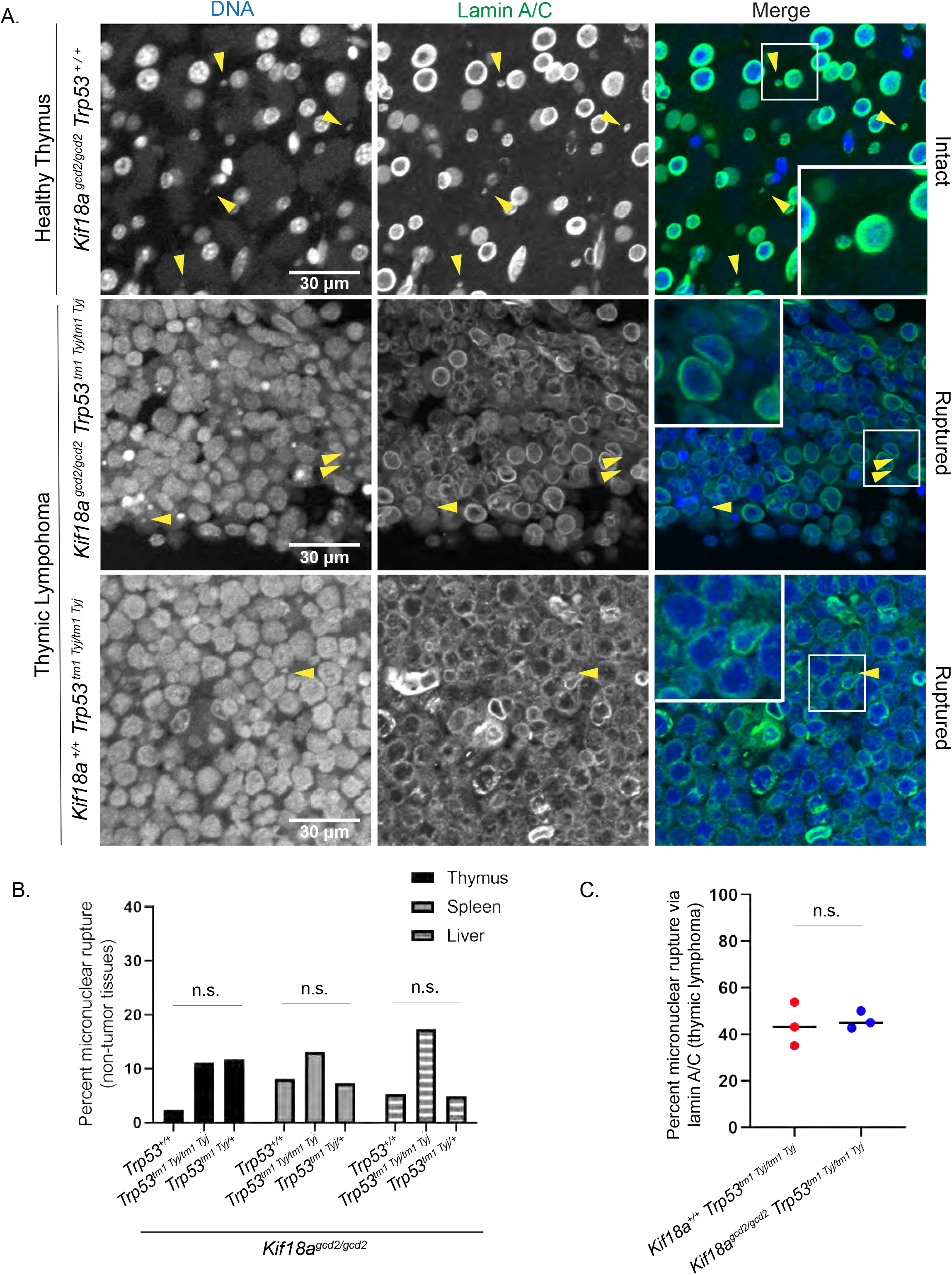
Micronuclear envelopes in *Kif18a* mutants are stable in healthy tissue but display increased rupture in tumor cells. **(A)** Representative images of nuclear envelopes for primary nuclei and micronuclei occurring in healthy thymus tissue (top) and in thymic lymphoma (middle and bottom) from the indicated genotypes. Sections were stained with Hoechst (DNA, blue) and lamin A/C (green); yellow arrowheads indicate micronuclei. **(B)** Plot showing the percent of micronucleated cells experiencing micronuclear envelope rupture in healthy thymus, liver, and spleen tissues, as determined via absence of lamin A/C signal co-occurring with micronuclear DNA. n = 3 tissue types per one biological replicate for genotype. *Kif18a^gcd2/gcd2^, Trp53^+/+^* (2/83 ruptured micronuclei in thymus, 6/74 ruptured micronuclei in spleen, 3/57 ruptured micronuclei in liver); *Kif18a^gcd2/gcd2^, Trp53^+/tm1^ ^Tyj^* (14/120 thymus, 5/68 spleen, 2/41 liver); *Kif18a^gcd2/gcd2^, Trp53^tm1^ ^Tyj/tm1^ ^Tyj^* (4/36 thymus, 16/122 spleen, 8/46 liver). Thymus, p = 0.052, n.s.; spleen p = 0.54, n.s.; liver p = 0.056, n.s. (Table S1.) Indicated p values were calculated by χ2 analysis. **(C)** Plot showing percent of micronuclei with ruptured nuclear envelopes in thymic lymphoma tissues from the indicated genotypes n=3 biological replicates per genotype: *Kif18a^gcd2/gcd2^, Trp53^tm1^ ^Tyj/tm1^* ^Tyj^ totaled 117/253 ruptured micronuclei from 4210 cells; *Kif18a^+/+^, Trp53^tm1^ ^Tyj/tm1^* ^Tyj^ 46/107 ruptured micronuclei from 3099 cells; p = 0.7648, n.s. (Table S3). Individual data points indicate individual biological replicates. Indicated p values were calculated via unpaired Student’s t-test.

We also investigated whether micronuclei within tumor tissues maintain stable nuclear envelopes. Sections from primary thymic lymphomas stained for lamin A/C and DNA (Hoescht) were used to analyze micronuclear rupture (Fig. 3A). Micronuclear envelope rupture rates in tumors were elevated relative to normal thymus tissue, however, there was no significant difference in the rupture frequency between mice lacking only p53 and those lacking both *Kif18a* and p53 (43% vs 46%, Fig. 3C and Table S3). These data suggest that the stability of micronuclear envelopes in *Kif18a* mutant cells could limit genomic instability and spontaneous tumorigenesis in *Kif18a* mutant mice.

### Micronuclei induced through loss of KIF18A rupture infrequently *in vitro*

To further explore micronuclear envelope stability in KIF18A mutant cells, we established an *in vitro* system to compare micronuclei induced via different types of insults in a human retinal pigment epithelial cell line (RPE1) immortalized by human telomerase expression (hTERT). hTERT-RPE1 cells are female, near diploid cells containing a modal chromosome number of 46 with a single derivative X chromosome and have been used previously for investigating micronuclear envelope rupture (Zhang, et al., 2015; Hatch et al., 2013; Liu et al., 2018). We will refer to these hTERT-RPE1 cells as “RPE1” throughout, for simplicity.

Micronuclei were induced in RPE1 cells via: 1) a nocodazole drug washout treatment, which leads to improper attachments between kinetochores and microtubules; 2) knockout (KO) of the *KIF18A* gene; 3) sub-lethal doses of radiation, which lead to double-stranded DNA breaks and fragmented chromosomes; and 4) siRNA knockdown of Mad2 (mitotic arrest deficient-2) protein, which disables the mitotic spindle assembly checkpoint and causes micronuclei through a combination of both improper kinetochore-microtubule attachments and chromosome unalignment (Fenech and Morley, 1985; Cimini et al., 2001; Burds, et al., 2005; Lusiyanti et al., 2016; Fonseca et al., 2019; Fig. 4A). Micronuclei also spontaneously form within wild type populations of RPE1 cells at low frequencies (1%), and RPE1 cells treated with a non-targeting siRNAs were used as controls (Tolbert et al., 1992).

**Figure 4:**
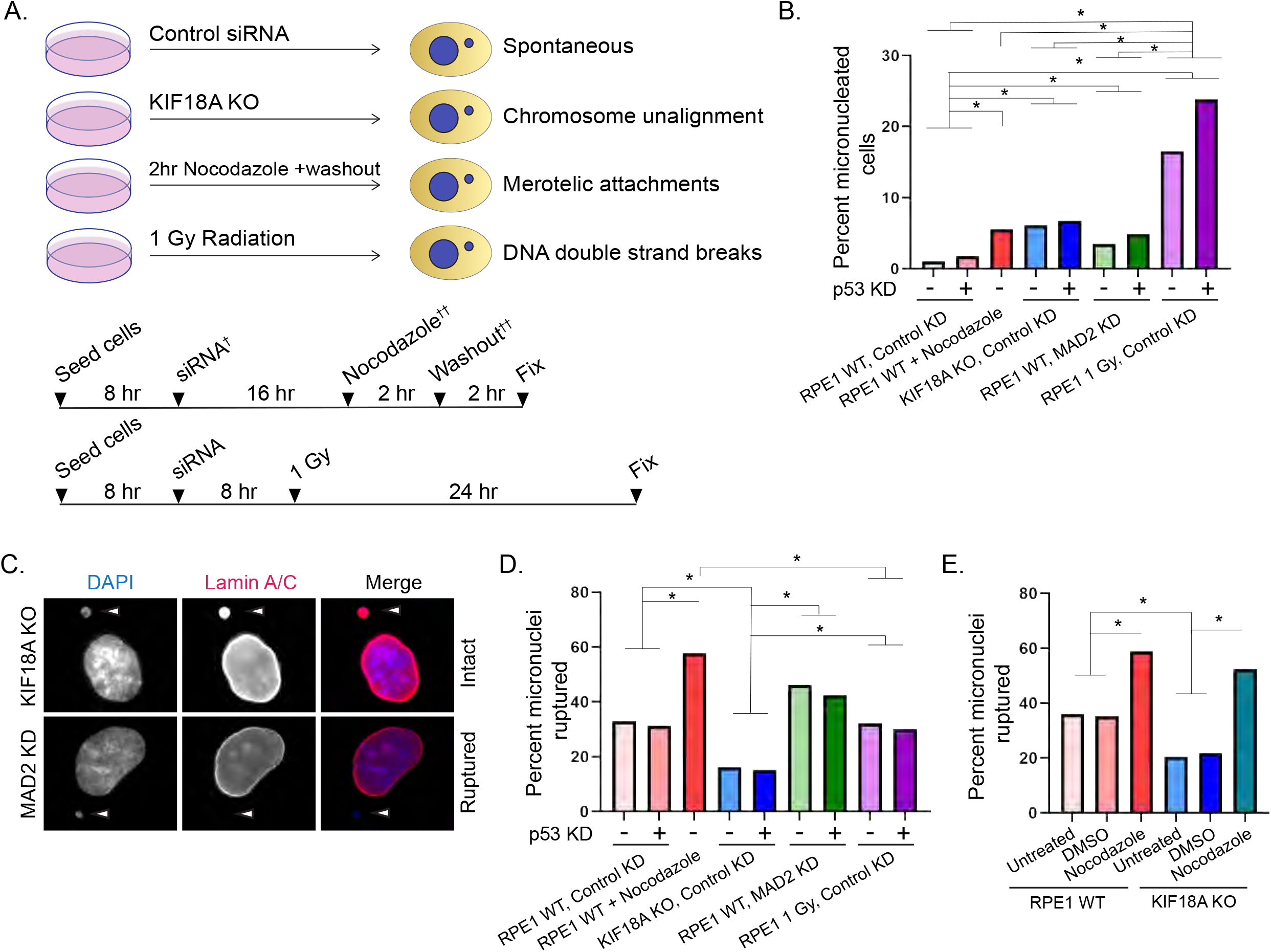
Micronuclei resulting from loss of KIF18A function in human cells rupture infrequently. **(A)** Schematic of experimental design. **(B)** Plot showing percent of micronucleated RPE1 cells following treatment with the indicated siRNAs or drug washout. Data are pooled from three independent experiments. n=4188 (RPE1 Control KD), n=3536 (RPE1 Control+p53 KD), n=661 (KIF18A KO Control KD), n=869 (KIF18A KO Control KD+p53 KD) n=1223 (RPE1 MAD2 KD), n=1157 (MAD2 KD+p53 KD), n=4005 (RPE1+nocodazole washout), n=2189 (RPE1 1 Gy, Control KD), n=3080 (RPE1 1 Gy, Control KD+p53 KD). *, p < 0.0001. (Table S4) **(C)** Representative images of fixed, micronucleated RPE1 cells labeled with DAPI and lamin A/C to visualize DNA and nuclear envelope, respectively. **(D)** Plot showing percentage of micronucleated cells that exhibited micronuclear envelope rupture in RPE1 cells, as determined by absence of lamin A/C signal, following the indicated treatments. Data are pooled from three independent experiments. n=485 (RPE1 Control KD), n=510 (RPE1 Control+p53 KD), n=807 (KIF18A KO Control KD), n=720 (KIF18A KO Control KD+p53 KD), n=631 (RPE1 MAD2 KD), n=648 (RPE1 MAD2 KD+p53 KD), n=726 (RPE1+nocodazole washout), n=622 (RPE1 1Gy, Control KD), n=778 (RPE1 1Gy, Control KD+p53 KD) *, p < 0.01. (Table S5.) **(E)** Plot showing percentage of micronuclei that exhibited micronuclear envelope rupture in RPE1 Control and KIF18A KO cells subjected to DMSO treatment or nocodazole-washout, as indicated. n=161 (RPE1 Untreated), n=162 (RPE1+DMSO washout), n=171 (RPE1+nocodazole washout), n=253 (KIF18A KO Untreated), n=293 (KIF18A KO+DMSO washout), n=278 (KIF18A KO+nocodazole wasout), * p < 0.01. (Table S6.) Data are from three independent experiments (B,C,E) and from four experiments (D). Indicated p values were calculated by χ2 analysis.

We analyzed cells following each treatment for the presence of micronuclei via staining with the DNA dye DAPI. Micronuclei were identified as DAPI-stained chromatin masses outside the main nucleus, and the percentage of micronucleated cells observed in each population was quantified. Consistent with previous observations, we found that 5.3% of KIF18A KO RPE1 cells formed micronuclei (Fig. 4B and Table S4) (Fonseca et al., 2019). To facilitate comparison, a short treatment of nocodazole (2 hours) before washout was used to yield a similar percentage of micronucleated cells (5.6%, Fig. 4B and Table S4). We also found that 4% of RPE1 cells treated with MAD2 siRNAs formed micronuclei and 19% of RPE1 cells subjected to 1 Gy radiation had formed micronuclei when evaluated 24-hours after exposure (Fig. 4B and Table S4).

Micronuclear envelope rupture was assessed by analyzing RPE1 cells labeled with lamin A/C antibodies and DAPI. Micronuclei were scored as ruptured if lamin A/C label was absent (Fig. 4C). To validate this approach, we tested the ability of micronuclei to retain an mCherry-tagged nuclear localization sequence (mCherry-N7) as a function of lamin A/C staining (Fig. S1). Specifically, micronuclear envelopes were quantified as ruptured based on 1) discontinuous lamin A/C staining and 2) leakage of mCherry-N7 signal outside the contained micronuclear area, identified via DAPI. By comparing these two criteria in the same micronuclei, we determined that lamin A/C signal alone is predictive of the integrity of the micronuclear envelope for 96% of cells (134/140 micronuclei), suggesting that lamin A/C immunofluorescence is a reliable method to determine rupture status of micronuclear envelopes in fixed cells. Consistent with previous reports (Hatch et al., 2013, and Liu et al., 2018), micronuclei produced via nocodazole washout experienced high rates of micronuclear envelope rupture (58%), as evidenced by a loss of robust lamin A/C signal co-occurring with micronuclear DNA (Fig. 4D and Table S5). In contrast, micronuclei in KIF18A KO cells exhibited low rates of micronuclear envelope rupture (16%) compared to micronuclei produced via all other inductions (p < 0.001, Fig. 4D and Table S5). We observed a moderate level of micronuclear envelope rupture in cells following MAD2 KD (46%), control KD (33%), and irradiation (32%, Fig. 4D and Table S5). It should be noted that micronuclei that form spontaneously (control KD) and those that form following MAD2 KD could result from a mix of initial cellular insults, including improper kinetochore microtubule attachments and alignment defects. This could explain the intermediate level of rupture observed compared to micronuclei in nocodazole treated and KIF18A KO cells. The frequencies of micronuclear envelope rupture in each population were not significantly affected by p53 KD, consistent with our *in vivo* results.

The relatively low micronuclear envelope rupture frequency observed in KIF18A KO cells could be explained by increased micronuclear envelope stability or a requirement for KIF18A in the micronuclear envelope rupture process. To distinguish between these possibilities, KIF18A KO and control RPE1 cells were subjected to nocodazole washout, in parallel, and micronuclear envelope integrity was assessed. Because cells were fixed 2 hours post nocodazole washout, a large fraction of micronuclei present in each cell population at this time point are expected to have formed due to improper kinetochore-microtubule attachments. We found that micronuclei in KIF18A KO cells treated with nocodazole washout displayed similar rates of rupture as those produced via drug treatment in RPE1 control cells (not significant, p = 0.2041, Fig. 4E and Table S6). These data indicate that KIF18A is not required for micronuclear envelope rupture, and therefore, micronuclear envelopes in KIF18A KO cells are more stable than those formed due to induced kinetochore microtubule attachment defects.

### Micronuclei in KIF18A KO cells successfully recruit non-core nuclear envelope components

Nuclear envelope stability is dependent on proper reassembly of numerous nuclear envelope components at the completion of cell division. Before chromosomes can interact with spindle microtubules in mammalian cells, nuclear envelope components must be disassembled and relocated. Several nuclear envelope proteins are found ubiquitously throughout the cytoplasm following nuclear envelope disassembly, whereas other components are unevenly distributed to organelles in the dividing cell (Hetzer, 2010). Some nuclear pore components are localized to the kinetochore (ex. ELYS, Nup107-160 complex), while inner nuclear membrane proteins, such as lamin B, are stored within the membranes of the ER at this time (Yang et al., 1997; Zuccolo et al., 2007; Hetzer, 2010).

In addition to differences in their mitotic localization, nuclear envelope components are categorized by their localization on decondensing chromatin surfaces following chromosome segregation (Clever et al., 2013; Liu and Pellman, 2019). Lamin A/C, a nuclear envelope protein found ubiquitously throughout the cytoplasm during mitosis, is a “core” nuclear envelope component, as it is recruited to the central chromosome mass nearest the central spindle axis during nuclear envelope reformation (Clever et al., 2013). Alternatively, lamin B is a “non-core” component because it is targeted to the chromosome peripheral regions during nuclear envelope reformation (Clever et al., 2013). Incorporation of non-core components is necessary for transport-competent nuclear envelopes and proper nuclear functions (Hetzer, 2010; Clever et al., 2013, Liu and Pellman, 2019). Micronuclear envelope stability is enhanced by successful recruitment of lamin B, while loss of lamin B causes holes to form in the lamina, increasing the frequency of nuclear envelope rupture (Vergnes et al., 2004; Vargas et al., 2012; Hatch et al., 2013; Liu et al., 2018).

To compare the extent of non-core nuclear envelope component recruitment to micronuclei in KIF18A KO cells and those subjected to nocodazole drug washout, cells were fixed and co-stained with DAPI and an antibody against lamin B1 (Fig. 5A). These experimental conditions were chosen for comparison because they exhibited the lowest and highest rates of micronuclear envelope rupture, respectively (Fig. 4D and Table S5). Fluorescence intensities of lamin B were measured using background-subtracted radial profile plots. The DNA boundary, as determined by DAPI, was used to determine the appropriate boundary of chromatin area for each individual micronucleus quantified. Ratios of lamin B surrounding micronuclei and corresponding primary nuclei within the same cell were also calculated for comparison. Quantifications of lamin B recruitment are reported both across populations of micronuclei, independent of rupture status (Fig. 5B-C), and as a function of micronuclear envelope integrity determined via lamin A/C (Fig. 5D). Lamin B levels in KIF18A KO cell micronuclei were significantly higher than those in nocodazole washout-treated cells and similar to those measured in primary nuclei (Fig. 5B-C; p < 0.01). These data indicate that lamin B recruitment to micronuclei is more efficient in KIF18A KO cells than in nocodazole treated cells.

**Figure 5:**
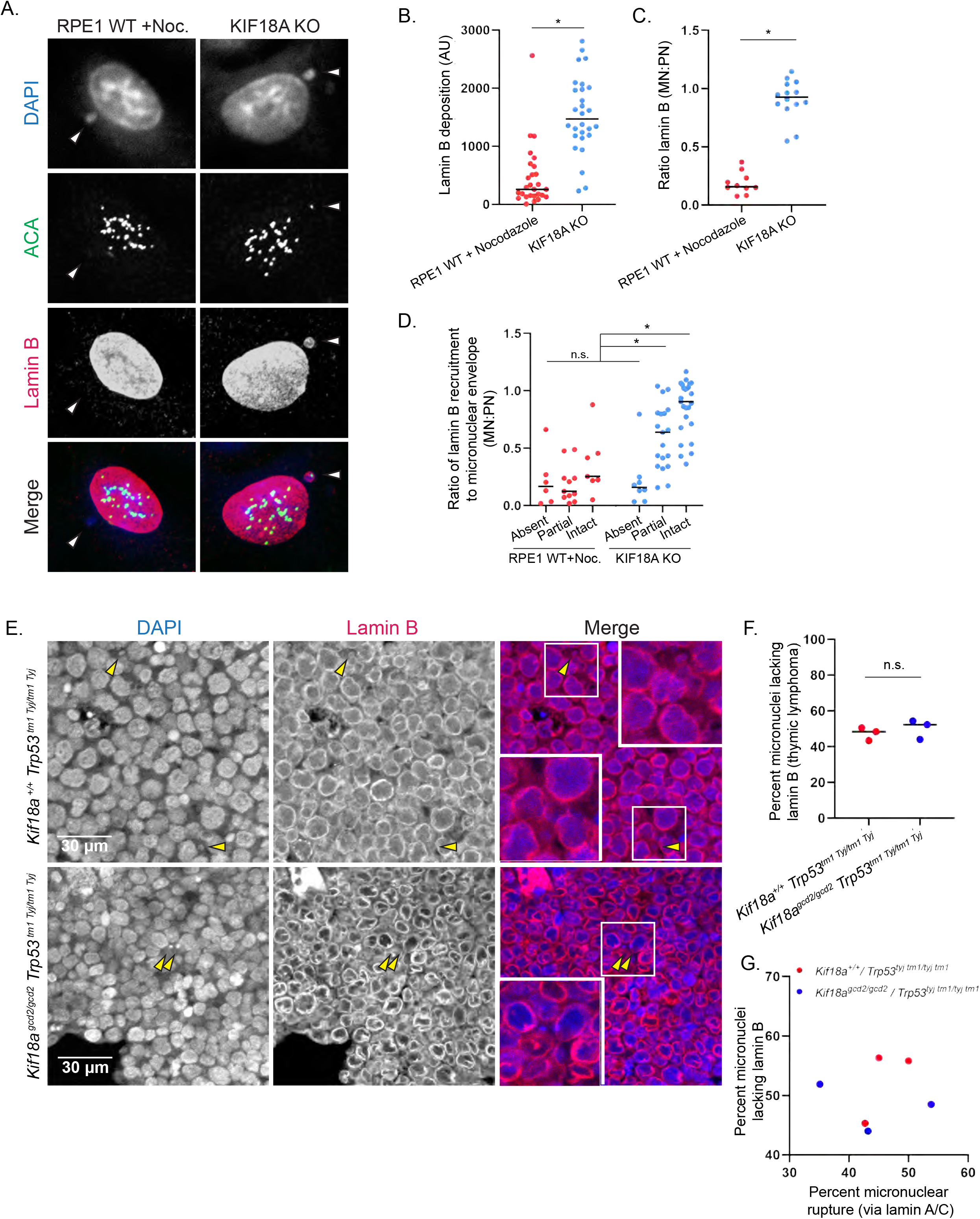
Micronuclei in KIF18A-deficient cells successfully recruit the non-core nuclear envelope component lamin B. **(A)** Representative images of fixed, micronucleated RPE1 cells labeled with DAPI, ACA, and lamin B to visualize DNA, centromeres, and a non-core nuclear envelope component, respectively. **(B)** Plot displaying lamin B fluorescence in micronuclear envelopes. Data are from three independent experiments; n=27 (RPE1+nocodazole washout) n=28 (KIF18A KO) * p < 0.001. Data points indicate individual micronuclei. **(C)** Plot displaying the ratio of lamin B recruited to the micronuclear (MN) envelope, directly compared to lamin B recruited to the primary nuclear (PN) envelope in the same cell. Data are from three independent experiments; n=10 (RPE1+nocodazole washout) n=14 (KIF18A KO) * p < 0.0001. Data points indicate individual micronucleated cells. **(D)** Plot displaying the ratios of lamin B recruited to the MN envelope compared to lamin B recruited to the PN envelope, parsed by MN integrity, as assessed via co-staining with lamin A/C antibody. n=24 (RPE1+nocodazole washout), n=52 (KIF18A KO); * p < 0.01. Data are from three independent experiments. Data points indicate individual micronucleated cells. **(E)** Representative images of thymic lymphoma tumor sections stained with Hoechst (DNA, blue) and lamin B (nuclear envelope, red) (micronuclei indicated via yellow arrowheads). **(F)** Plot showing percentage of micronuclei in thymic lymphoma tissues that lacked lamin B. n=3 biological replicates per genotype were assessed, totaling: *Kif18a^+/+^, Trp53^tm1^ ^Tyj/tm1^* ^Tyj^ 150/311 micronuclei lacking lamin B; *Kif18a^gcd2/gcd2^, Trp53^tm1^ ^Tyj/tm1^* ^Tyj^ 200/387 micronuclei lacking lamin B; p = 0.5048, n.s. (Table S7). Data points indicate individual biological replicates. (G) Plot showing comparison of each sample’s percentage of micronuclei in thymic lymphoma tissues that lacked lamin B, plotted by percentage of ruptured micronuclei (indicated by loss of lamin A/C), Indicated p values for numerical data were obtained via unpaired Student’s t-test, for comparisons among two conditions, or a one-way ANOVA with post-hoc Tukey test, for comparisons among more than two conditions.

To determine if lamin B is also recruited to micronuclei in *Kif18a* mutant cells *in vivo*, we investigated lamin B in thymic lymphoma tissues co-stained with Hoechst (DNA, Fig. 5E). We found similar levels of lamin B recruitment to micronuclei within *Kif18a^+/+^, Trp53^tm1^ ^Tyj/tm1^* ^Tyj^ and *Kif18a^gcd2/gcd2^, Trp53^tm1^ ^Tyj/tm1^* ^Tyj^ tissues, where 48% and 52% of micronuclear envelopes, respectively, lack robust recruitment of lamin B (Fig. 5F and Table S7). These results are consistent with the level of rupture measured via lamin A/C staining and indicate that lamin B is recruited to stable micronuclei within *Kif18a* mutant mice (Fig. 3B and 5F). A comparison of micronuclear membranes from the same tissues (but not the same cells) supports the idea that lamin B recruitment correlates with rupture frequency (Fig. 5G).

### Micronuclei in KIF18A KO cells exhibit successful nuclear envelope expansion

Nuclear envelope stability also depends on efficient recruitment of membrane and membrane components from the ER as chromosomes decondense and nuclear area expands in late mitosis (Anderson and Hetzer, 2008; Hetzer, 2010; Clever et al., 2013; De Magistris and Antonin, 2018). Multiple nuclear envelope and nuclear pore components have been found to directly impact chromatin decondensation and chromatin remodeling, upon being targeted to chromatin (Chi et al., 2007; Korfali et al., 2010; Kuhn, et al., 2019). Data from intact cells, as well as *in vitro* nuclear assembly systems, show that physically disrupting the connection between nuclei and the peripheral ER (where nuclear envelope components are stored during mitosis) can indeed block nuclear expansion (Anderson and Hetzer, 2007). To investigate whether micronuclei in KIF18A KO cells are able to successfully expand and stabilize, we measured the change in both micronuclear and primary nuclear chromatin area over time as chromosomes decondensed in live telophase cells (Fig. 6A). Micronuclear chromatin within KIF18A KO cells exhibited a 1.4-fold increase in area, similar to that of chromatin within primary nuclei (Fig. 6B-D). In contrast, micronuclei forming as a result of nocodazole washout exhibited a significantly reduced expansion during telophase (p < 0.01). These results suggest that micronuclei forming after nocodazole washout experience chromatin restriction, which may increase the frequency of micronuclear envelope rupture.

**Figure 6:**
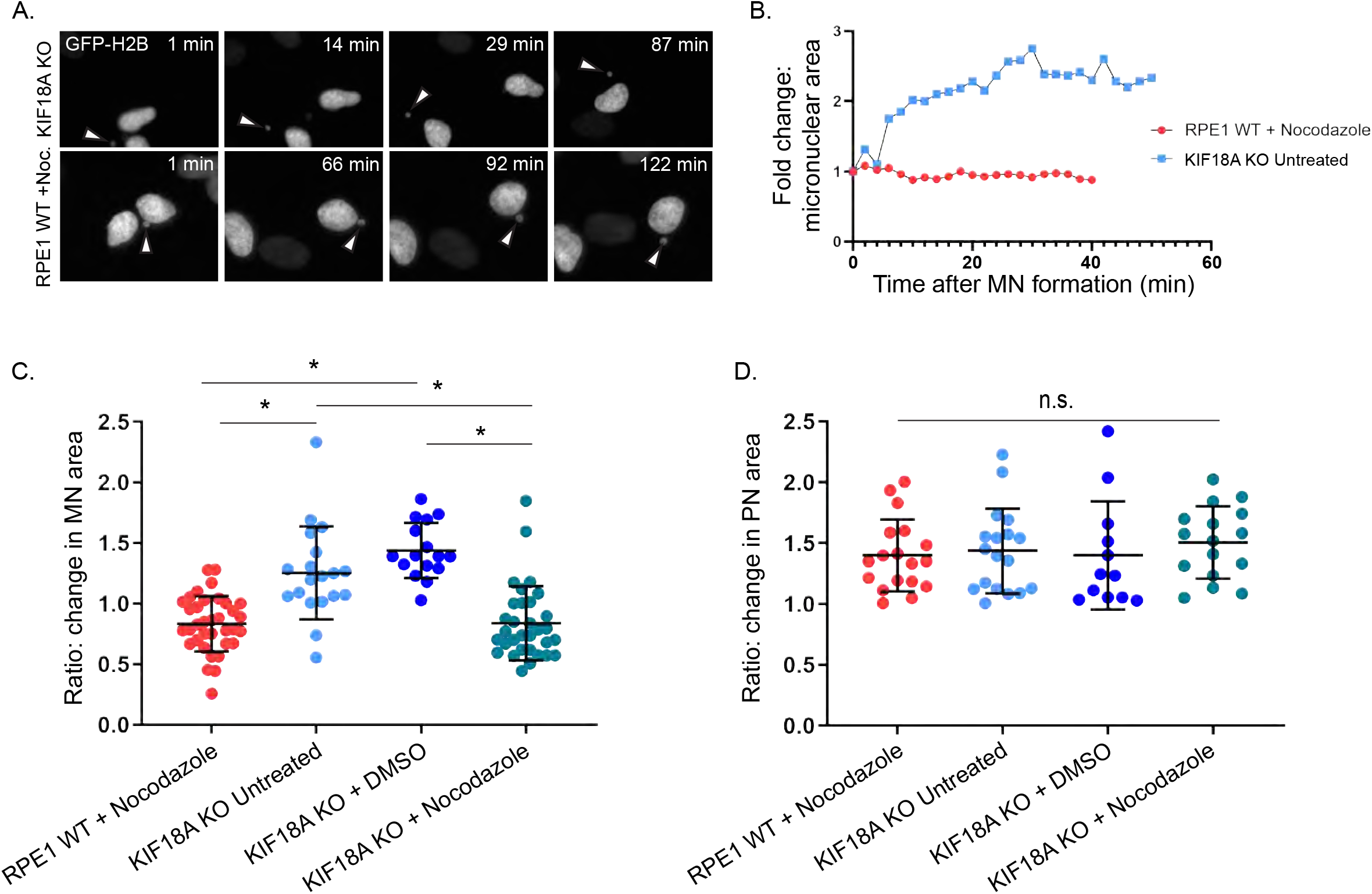
Micronuclei in KIF18A KO cells exhibit chromatin expansion upon exit from mitosis. **(A)** Stills from time-lapse imaging of micronuclei (indicated via arrowhead) arising due to unalignment in a KIF18A KO RPE1 cell, top; and in an RPE1 control cell following nocodazole washout treatment, bottom. Cells were transfected with GFP-H2B to label DNA. **(B)** Representative traces displaying fold change in micronuclear chromatin area beginning immediately after completion of chromosome segregation at the time of initial micronucleus formation until chromatin was decondensed. Traces shown in B match representative images shown in A. Individual fold change trace indicates a single representative micronucleus per condition. **(C)** Plot of final fold change in micronuclear area (final area divided by initial recorded micronuclear area), for the indicated conditions. Data points represent individual micronuclei. n=35 (RPE1+nocodazole), n=19 (KIF18A KO untreated), n=16 (KIF18A KO + DMSO), n=32 (KIF18A KO+nocodazole). Data were collected from four independent experiments, * p < 0.0001). Data points indicate individual micronuclei. Error bars indicate standard deviation about the mean. **(D)** Final ratio of fold change in primary nuclear area, from the same cells that micronuclei were measured in C. n=18 (RPE1+nocodazole), n=18 (KIF18A KO untreated), n=12 (KIF18A KO+DMSO), n=16 (KIF18A KO+nocodazole); p = 0.80, n.s. Data points indicate individual daughter primary nuclei. Error bars indicate standard deviation about the mean. Statistical comparisons were made using a one-way ANOVA with Tukey’s multiple comparisons test.

### Lagging chromosomes in KIF18A KO cells are found near main chromatin masses

Previous work in both cultured human cells and *Drosophila* has demonstrated that micronuclear envelope stability is influenced by the subcellular location of nuclear envelope assembly around individual lagging chromosomes within the mitotic spindle (Afonso et al., 2014 and Maiato et al., 2015, Liu et al., 2018). Thus, we compared the locations of lagging chromosomes in KIF18A KO and nocodazole washout RPE1 cells. Asynchronously dividing *KIF18A* KO cells were fixed and labeled with an antibody against γ-tubulin, to mark spindle poles, and DAPI, to stain chromatin. Cells in late anaphase were scored for the presence of late-lagging chromosomes, which notably trailed behind the main chromatin masses. Late lagging chromosomes were observed in 44% (52/118) of nocodazole-washout treated RPE1 cells, and in 9% (4/43) of *KIF18A* KO cells, respectively (Fig. S2A-B; p < 0.001). These data agree with our prior conclusion that chromosomes in the midzone during late anaphase are rare in KIF18A KO cells and suggest that differences in lagging chromosome positions may underscore the differences in micronuclear envelope stability exhibited by KIF18A KO and nocodazole washout cells (Fonseca et al., 2019).

To explore this question further and obtain precise measurements of lagging chromosome positions from a larger number of KIF18A KO cells, we treated with a CDK-1 inhibitor to enrich cells in G2 and then released into mitosis by drug washout. This facilitated a similar enrichment of late anaphase cells as observed following nocodazole washout. Cells were then fixed and stained for antibodies against γ-tubulin and centromeres (Fig. 7A). The positions of individual chromosomes relative to the spindle pole were measured in each half-spindle of anaphase cells. Centromere signals located further than two standard deviations from the average centromere position within each half spindle were identified as “lagging” (Fig. 7B). Lagging chromosomes in KIF18A KO cells were located closer to the main chromatin masses than those in nocodazole treated cells (Fig. 7C). In addition, the angle of each late-lagging chromosome centromere relative to the pole-to-pole axis was measured. There was not a significant difference between the angles of lagging chromosomes to the spindle in the two conditions (Fig. 7D). These data indicate that lagging chromosomes in nocodazole washout cells are closer to the midzone than those in KIF18A KO cells, which could contribute to the observed decrease in micronucleus stability measured in nocodazole treated cells.

**Figure 7:**
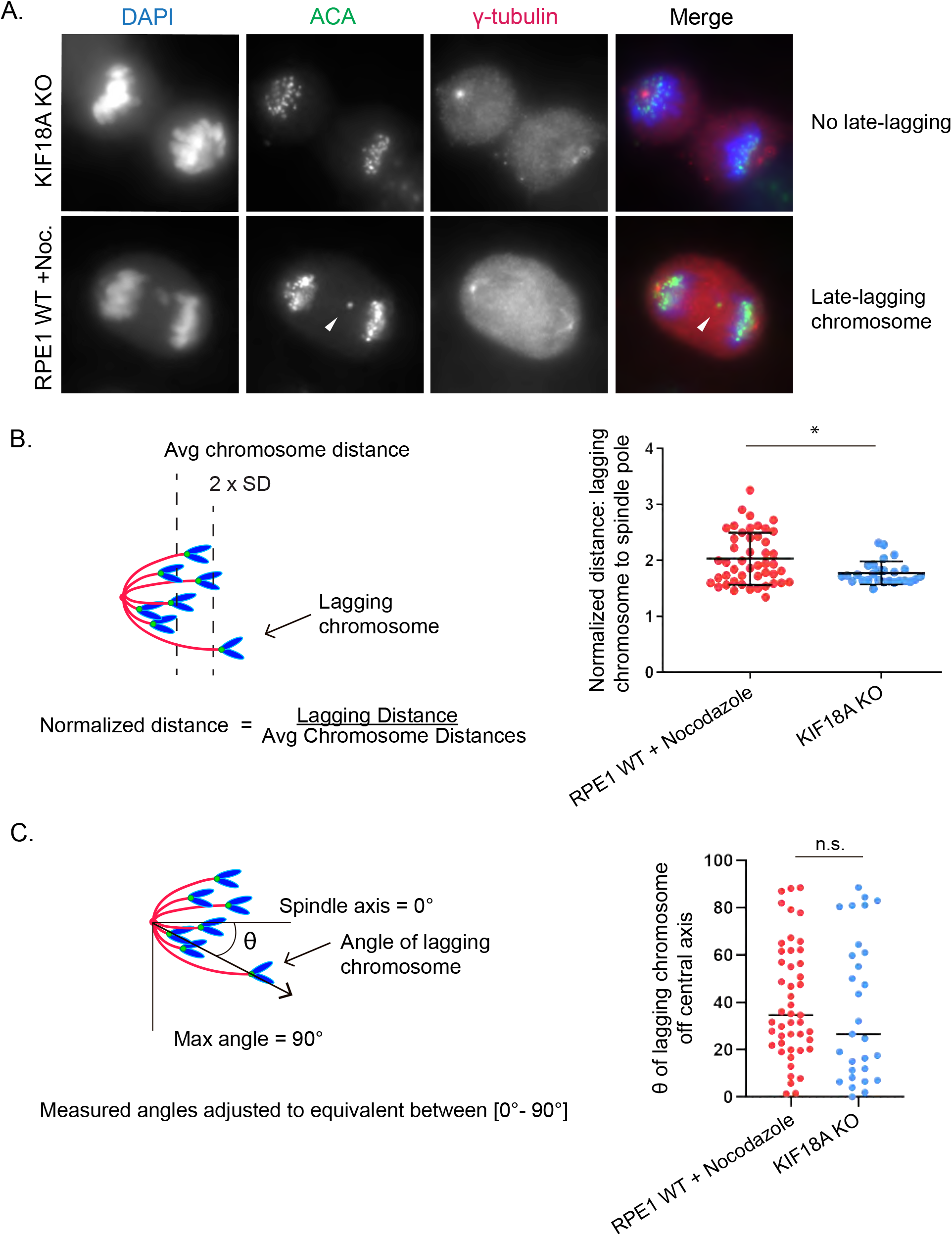
Lagging chromosomes in KIF18A KO cells are located near the spindle poles in late anaphase. **(A)** Representative images of late anaphase RPE1 cells that were fixed and labeled with antibodies against centromeres and spindle poles. Arrowhead indicaes ‘lagging’ chromosome. **(B)** Left: Schematic depicting how lagging chromosome positions were measured and normalized within each half-spindle. Right: Plot showing normalized late lagging chromosome to spindle pole distances measured in KIF18A KO RPE1 cells and nocodazole-washout treated RPE1 WT cells. n=47 (RPE1 WT+nocodazole), n=27 (KIF18A KO). Data were collected from three independent experiments, * p < 0.01. Data points indicate individual lagging chromosomes. Error bars represent standard deviation about the mean. **(C)** Left: Schematic depicting how lagging chromosome angles were measured relative to the central spindle axis. Measured angles were converted to equivalent angles within the range of 0 and 90 degrees. Right: Plot of late lagging chromosome angles relative to the central spindle axis for the indicated conditions (n = 29, KIF18A KO and n = 47, RPE1 nocodazole-washout; p = 0.186, n.s.). Data were collected from three independent experiments. Data points indicate individual lagging chromosomes. Statistical comparisons were made using a two-tailed unpaired Student’s t-test.

To explore other possible explanations for the differences in micronuclear envelope stability in KIF18A KO and nocodazole washout cells, we investigated micronuclear chromatin area, the DNA damage marker γH_2_AX, and presence of centromeres. However, consistent with previous reports, none of these factors had strong correlations with micronuclear envelope stability (Fig. S3, A-D; Hatch et al, 2013).

## Discussion

Micronuclei have been proposed not only as passive markers of genomic instability, but also as active drivers of tumorigenesis (Stephens et al, 2011; Rausch et al., 2012; Holland and Cleveland, 2012; Crasta et al., 2012; Nones et al., 2014; Zhang et al., 2015; Luijten et al., 2018). Inconsistent with this proposal, *Kif18a* mutant mice readily form micronuclei *in vivo* but do not spontaneously develop tumors. We investigated this apparent contradiction by testing the contributions of two, non-mutually exclusive models: (1) that p53 activity in *Kif18a* mutant mice prevents propagation of micronucleated cells and the subsequent reincorporation of damaged DNA into primary nuclei and (2) that micronuclei form stable nuclear envelopes. Our data favor the second model, and indicate that micronuclei in *Kif18a* mutant cells, which form as a result of mitotic chromosome alignment defects (Fonseca et al, 2019), have stable nuclear envelopes that undergo expansion as cells exit mitosis. The more stable micronuclear envelopes in KIF18A KO cells are less prone to experiencing rupture events, as assessed both in cultured cells and *in vivo* tissue sections, and these behaviors are independent of p53 status. In contrast, micronuclei that resulted from improper kinetochore microtubule attachments induced by nocodazole washout formed further from mitotic spindle poles, had unstable nuclear envelopes, and failed to undergo expansion (Hatch et al., 2013, Zhang et al., 2015, Liu et al., 2018). Taken together, this work demonstrates that the underlying cause of lagging chromosomes can strongly impact the stability of micronuclear envelopes that form around them, and therefore, their threat to genomic stability (Ding et al., 2003; Hoffelder et al., 2004; Terradas et al., 2009; Terradas et al., 2010; Huang et al., 2011, Crasta et al., 2012, Hatch et al., 2013, Zhang et al., 2015, Liu et al., 2018).

Loss of Kif18a had modest or no effect on survival of *Trp53* homozygotes and heterozygotes, respectively. These data are inconsistent with the idea that p53-dependent cell cycle arrest prevents micronuclei from promoting tumor development in *Kif18a* mutant mice and also indicate that, unlike colitis-associated colorectal tumors, *Kif18a* is not required for the growth of thymic lymphoma (Zhu et al., 2013). However, we cannot formally rule out that differences in tumor spectrum between *Kif18a^gcd2/gcd2^* , *Trp53^tm^ ^1^ ^Tyj/tm^ ^1^ ^Tyj^* and *Kif18a^gcd2/gcd2^* , *Trp53^tm^ ^1^ ^Tyj/+^* genotypes could also contribute to their differences in survival. In addition, our results suggest that the formation of micronuclei per se does not necessarily lead to tumorigenesis. This, together with prior studies which (1) detect no apparent increase in aneuploidy in *Kif18a^gcd2/gcd2^* mouse embryonic fibroblasts, and (2) demonstrate that *Kif18a* mutant mice are not predisposed to tumor formation when challenged with colitis-associated colorectal cancer, suggests that *Kif18a* mutant mice are relatively genomically stable despite elevated levels of detectable micronuclei in peripheral blood and tissues (Czechanski et al., 2015; Zhu et al., 2013). Thus, all micronuclei may not have the same potential to alter genomic stability, and additional physical or genetic insults are likely required to create permissive environments for micronuclei to drive cellular transformation in the cells or tissues where they form.

Our data indicate that the positioning of late-lagging chromosomes, which often form micronuclei, is impacted by the type of insult leading to the chromosome segregation error. Prior studies that characterized the impact of micronuclei on genomic stability primarily utilized treatments, such as nocodazole washout, that promote improper attachments between kinetochores and microtubules and give rise to micronuclei that form in the central-spindle, far from the spindle poles (Crasta et al., 2012, Zhang et al., 2015, Hatch et al., 2015, Liu et al., 2018). In contrast, lagging chromosomes in KIF18A KO cells were located near the main chromatin masses in late anaphase. The fact that micronuclei in KIF18A KO cells are relatively stable compared to those in nocodazole treated cells is consistent with work indicating that bundled microtubules and a gradient of Aurora B inhibit proper nuclear envelope reformation near the center of anaphase spindles (Afonso et al., 2014 and Maiato et al., 2015, Liu et al., 2018). Our data also indicate that the distance of a lagging chromosome from the pole is more important for nuclear envelope reformation than its position relative to the pole-to-pole axis.

On the other hand, chromosome size, prevalence of DNA damage, and whether the micronuclear chromatin contained centromeres did not strongly correlate with micronuclear envelope rupture status, consistent with previous work (Hatch et al., 2014). While we cannot rule out that there are other factors contributing to observed differences in micronuclear envelope stability, our data are consistent with the idea that lagging chromosome position strongly influences nuclear envelope stability and that this effect may be relevant *in vivo*.

In addition to the negative regulation proposed by microtubule bundling and inhibitory Aurora gradients, it is possible that positive regulation by the spindle poles and ER membranes may promote stable nuclear envelope reformation. This idea is consistent with our observations that pole-proximal positioning of lagging chromosomes correlates with successful expansion of micronuclear chromatin area and recruitment of lamin B to micronuclear membranes in KIF18A KO cells. For example, if lagging chromosomes are positioned nearer to the stores of nuclear envelope components located in the mitotic ER, this pole-proximal location may enhance prompt recruitment of necessary proteins and membrane to micronuclear envelopes during telophase. Potential positive regulators that impact nuclear envelope stability will require further investigation.

Why does micronuclear envelope stability differ between normal and tumor tissues? One possibility is that changes occurring during the process of cellular transformation may act to increase the frequency of micronuclear rupture. For example, a reduction of lamina has been associated with reduced stress resistance in the nuclear envelope, resulting in more frequent nuclear blebbing and rupture (Vergenes et al., 2004; Vargas et al., 2012; Hatch et al., 2013; Denais, et al., 2016). On the other hand, upregulation of lamina expression in transformed cancer cells enables nuclei to withstand greater mechanical forces and limit nuclear deformation, though this nuclear stiffness directly limits invasion efficacy (Vortmeyer-Krause et al., 2020). These conclusions suggest a tuned balance must be struck in the regulation of nuclear envelope dynamics for transformed cells to both withstand increased cytoskeletal forces and promote invasion efficiency (Vortmeyer-Krause et al., 2020). The increased stiffness of tumorous tissue architecture, paired with increased cytoskeletal forces present, may contribute to the increased prevalence of micronuclear envelope rupture in this context. Though growing evidence links increased mechanical stress to elevated rupture of primary nuclei in cancer cells, more research is needed to completely map these mechanistic changes to micronuclear envelope rupture in an *in vivo* context.

In conclusion, our work raises a number of interesting questions about the impact of micronuclei *in vivo.* To what extent and under what cellular conditions do micronuclei behave as drivers of tumorigenesis, and barring those conditions, are micronuclei simply passive biomarkers of instability? How does the surrounding tissue architecture impact transmitted cytoskeletal forces and alter micronuclear behaviors and rupture incidence? To what degree are immune system inflammatory sensors successful at recognizing and clearing damaged extranuclear DNA *in vivo,* and how might inflammatory micronucleation alter the progression of tumorigenesis at an organismal level? The development of new models that allow investigators to tune micronuclear rupture *in vivo* will be required to fully understand the impact of micronuclei.

## Materials and Methods

### Animal Ethics Statement

All procedures involving mice were approved by The Jackson Laboratory’s Institutional Animal Care and Use Committee and performed in accordance with the National Institutes of Health guidelines for the care and use of animals in research.

### Mouse development

A cohort of laboratory mice heterozygous for the null alleles, *Trp53^tm1Tyj^* and *Kif18a^gcd2^*, were generated by *in vitro* fertilization of oocytes from 30 heterozygous *Trp53*^Tm1TyJ^ females (JAX JR#2526, C.129S2(B6)-*Trp53*<tm1Tyj>/J) with sperm from a B6.*Kif18a*^gcd2 male (JAX JR#10508 / MMRRC #034325-JAX). A total of 143 offspring (64F/69M) were obtained from this expansion and those that were doubly heterozygous for each allele were intercrossed to produce a large cohort of animals for survival analysis. Cohort size and sample groups by genotype were based on published tumorigenesis and survival data on the *Trp53*<tm1Tyj>/ allele (PMID: 7922305) and z-tests, alpha = 0.5, 85% power. Genotypes and sample groups, excluding censored individuals) were as follows: *Heterozygous p53: Kif18a^gcd2/gcd2^*, *Trp53^tm1Tyj^*^/+^ (n = 40F, 35M) vs. *Kif18a*^+/+^, *Trp53^tm1Tyj^*^/+^ (40F, 38M). Homozygous p53: *Kif18a^gcd2/gcd2^*, *Trp53^tm1Tyj/tm1Tyj^* (n = 22F, 20M) vs. *Kif18a^+/+^*, *Trp53 ^tm1Tyj/tm1Tyj^* (n = 25F, 33M) or *Kif18a^gcd2/+^*, *Trp53 ^tm1Tyj/tm1Tyj^* (n = 13F, 20M). A cohort of *Kif18a^gcd2/gcd2^*, *Trp53^+/+^* (n = 38F, 34M) genetic background controls were included and no tumors were observed in this group of mice.

Two approaches were used for survival analysis. Prism 8 and R were used to generate survival curves and to perform Wang-Allison tests. These tests compare of the number of subjects alive and dead beyond a specified time point (90% percentile) between two sample groups. To account for censored data, and a nonparametric log-rank test as also used to compare the survival distributions. The log-rank test results in a Chi-square statistic, which was then used to calculate significance of the test.

### Peripheral blood micronucleus assays

Micronuclear assays of peripheral blood were conducted as previously described (Dertinger et al., 1996; Reinholdt et al., 2004). Peripheral blood was collected from the retro-orbital sinus of male and female laboratory mice, 12-18 weeks of age, for each of the following genotypes (4 males and 4 females per genotype). Genotypes were as follows: *Kif18a^gcd2/gcd2^*, *Trp53^tm1Tyj/tm1Tyj^ ; Kif18a^+/+^*, *Trp53 ^tm1Tyj/tm1Tyj^ Kif18a^gcd2/+^*, *Trp53 ^tm1Tyj/tm1Tyj^*; *Kif18a^+/+^*, *Trp53 ^tm1Tyj/tm1Tyj^ ; Kif18a^+/+^*, *Trp53^+/+^*. A cohort of *Kif18a^gcd2/gcd2^*, *Trp53^+/+^* (n = 38F, 34M). Briefly, 75 μl of blood was immediately mixed with 100 μl of heparin, and the mixture was then pipetted into 2 ml of ice-cold (−80°C) 100% methanol with vigorous agitation to prevent clumping. Samples were stored at −80°C overnight before processing for flow cytometry. *Sample preparation and flow cytometry:* Each blood sample was washed with 12 ml of sterile, ice-cold bicarbonate buffer (0.9% NaCl, 5.3 mM sodium bicarbonate, pH7.5), centrifuged at 500g for 5 min. and resuspended in a minimum of carryover buffer (~100 Dl). 20 Dl of each sample was added to a 5 ml polystyrene round-bottomed tube, and to each sample an 80 Dl solution of CD71-FITC and RNAseA (1 mg/ml) was added. Additional control samples were CD71-FITC alone and an additional sample with bicarbonate buffer alone to which propidium iodide (PI) would be later added (see below). Cells were incubated at 4C for 45 minutes, washed with 2 ml cold bicarbonate buffer, and centrifuged as above. Cell pellets were stored on ice and then, immediately prior to flow cytometric analysis, resuspended in 1 ml of ice-cold PI solution (1.25 mg/ml) to stain DNA. *Flow cytometry*: Samples were processed on a BD Bioscience LSRII fluorescence-activated cell sorter gated for FITC and PI, and set to collect 20,000 CD71 positive events at 5,000 events / sec. The CD71-FITC and PI control samples were used to calibrate for autofluorescence. Reticulocytes (Retic, CD71+, PI-[in the presence of RNAse A]), mature red blood cells (RBC, CD71 -, PI -), micronucleated normochromatic erythrocytes (NCE-MN, CD71-, PI+) and micronucleated reticulocytes (Ret-MN, CD71+, PI+) were measured using FlowJo software. The total % of spontaneous micronuclei in NCE was NCE-MN/(NCE-MN + RBC)*100.

### Cell Culture and transfections

hTERT-RPE1 cells (ATCC) were maintained at 37°C with 5% CO_2_ in MEM-α (Life Technologies) containing 10% FBS (Life Technologies) and 1% antibiotics. The Kif18A-deficient CRISPR line was produced as previously described (Fonseca et al, 2019). For fixed cell assays, cells were seeded on 12-mm acid-washed coverslips and transfected with 30 pmol siRNA complexed with RNAiMAX, following the manufacturer’s instructions. For live cell imaging, cells were subjected to plasmid transfections ([2 μg]/ each plasmid), performed using a Nucleofector 4D system, according to the manufacturer’s instructions (Lonza). RPE1 cells were transfected with SF solution and electroporated with code EN-150. Following electroporation, cells were seeded in 35-mm poly-L-lysine-coated glass bottom dishes (MatTek) 8 h before the addition of siRNA.

### Primary murine tumor extraction and histology sections

For derivation of primary murine tumors, mice were euthanized when they showed signs of labored breathing (suspected thymic lymphoma) or exhibited visible tumors causing difficulties in mobility (suspected muscular sarcoma). Mice were dissected, tissue samples were taken for genotype verification, and individual tumors were excised, washed in cold PBS, halved, and fixed in normal buffered formalin and embedded in paraffin. Five micrometer, step-sections were mounted on microscope slides and 10 slides per tissue sample were used for immunolabeling. Mounted paraffin sections were deparaffinized in xylene and rehydrated through a gradual ethanol series. Antigen retrieval was performed by boiling 0.01 M citric acid (pH 6) for 20 min. Slides were blocked in 20% goat serum in AbDil for one hour before antibody incubation overnight and staining via Hoechst. Stained sections were mounted in ProLong Gold without DAPI.

Tissue samples were genotyped by The Jackson Laboratory Transgenic Genotyping Service. Tissue sectioning and slide preparation was performed by The Jackson Laboratory Histolopathology Services.

### Plasmids and siRNAs

H2B-GFP was a gift from Geoff Wahl (The Salk Institute, La Jolla, CA; Addgene plasmid no. 11680). mCherry-Nucleus-7 plasmid was generated by Michael Davidson and obtained from Addgene (Addgene plasmid no. 55110). Cells were transfected using siRNAs targeting *MAD2* sequence 5′-AGAUGGAUAUAUGCCACGCTT-3′ (Qiagen), pools of siRNAs targeting the *TP53* sequence 5’-GAAAUUUGCGUGUGGAGUA-3’, 5’-GUGCAGCUGUGGGUUGAUU-3’, 5’-GCAGUCAGAUCCUAGCGUC-3’, and 5’-GGAGAAUAUUUCACCCUUC-3’ (Dharmacon ON-TARGETplus), or negative control *Silencer* siRNA #2 (ThermoFisher Scientific).

### Cell fixation and immunofluorescence

RPE1 cells were fixed in −20°C methanol (Thermo Fisher) and 1% paraformaldehyde (Electron Microscopy Sciences). Cells were then washed in 1x TBS and blocked in antibody dilution buffer (Abdil; tris buffered saline, pH 7.4, 1% bovine serum albumin, 0.1% Triton X-100, and 0.1% sodium azide) containing 20% goat serum. Cells were incubated with the following primary antibodies for 1 h at room temperature in Abdil: mouse anti-human lamin A/C (1:200; Millipore), rabbit anti-lamin A/C (1:200; Abcam ab26300), rabbit anti-lamin B1 (1:200; Abcam ab16048), rabbit anti-γ-H_2_AX (1:200; COMPANY), rabbit anti-mCherry (1:200; Abcam ab167453), mouse anti-γ-tubulin (1:200; Sigma-Aldrich). Cells were incubated overnight at 4°C with human anti-centromere antibody (ACA; 1:200; Antibodies Inc.). Cells were incubated for 1 h at room temperature, in the dark, with goat secondary antibodies against mouse, rabbit, or human IgG conjugated to Alex Fluor 499, 594, or 647 (Molecular Probes by Life Technologies). Coverslips were mounted on glass slides with Prolong Gold antifade reagent plus DAPI (Molecular Probes by Life Technologies).

### Microscopy

Cells were imaged on a Nikon Ti-E inverted microscope (Nikon Instruments) controlled by NIS Elements software (Nikon Instruments) with a Spectra-X light engine (Lumencor), Clara cooled-CCD camera (Andor), 37°C environmental chamber, and the following Nikon objectives: 20× Plan Apo differential interference contrast (DIC) M N2 (NA 0.75), 40× Plan Apo DIC M N2 (NA 0.95), 60× Plan Apo λ (NA 1.42), and 100× APO (NA 1.49).

Imaging of nuclear envelope component lamin B1 for assessment in RPE-1 cells and lamin A/C or lamin B1 assessment in histological sections of mouse tissues was performed at the Microscopy Imaging Center at the University of Vermont. Fixed cells or tissues were imaged using a Nikon A1R-ER confocal microscope (Nikon Instruments), controlled by NIS Elements software (Nikon Instruments), with a Sola light engine (Lumencor), Nikon A1plus camera, containing a hybrid resonant and high resolution Galvano galvanometer scanhead set on an inverted Ti-E system (Nikon Instruments), and the following Nikon objectives: 20× Plan Apo λ (NA 0.75), 40× Plan Fluor Oil DIC H N2 (NA 1.3), 60× APO TIRF Oil DIC N2 (NA 1.49).

### Live cell imaging

Cells were transferred into CO_2_-independent media with 10% FBS and 1% antibiotic (Life Technologies) for imaging via fluorescence microscopy. For long term imaging of H2B-GFP and/or mCherry-Nucleus7 expressing cells, single focal plane images were acquired at 2-minute intervals with a 20x or 40x objective. For imaging live micronucleus production and chromatin decondensation/expansion, fields were scanned and metaphase cells were selected for imaging.

### Micronuclear envelope rupture counts

For fixed cells, micronucleus counts were made using single focal plane images of DAPI stained cells. Image acquisition was started at a random site at the bottom edge of the coverslip, and images were acquired every two fields of view using a 20x objective for micronucleus frequencies in each population. Micronuclear envelope rupture frequencies were made using single focal plane images of DAPI stained cells. Image acquisition was started at a random site at the bottom of the coverslip, all micronuclei that were observed were imaged using a 20x objective, until ~50 micronuclei were found via DAPI stain only (blind to nuclear envelope marker) in each condition.

To validate the use of lamin A/C presence as a marker of intact micronuclear envelopes, RPE1 cells were transfected with mCherry-N7 plasmid (mCherry tag fused to nuclear localization sequence repeats to allow targeted nuclear import of plasmid), seeded onto 12-mm acid-washed coverslips, and maintained for 24 h. Cells were fixed and stained with mouse anti-lamin A/C and rabbit-anti-mCherry, and mounted on glass slides with Prolong Gold antifade reagent plus DAPI. Coverslips were scanned in the DAPI channel only, and all micronuclei found (up to 50 micronuclei per experiment) were imaged in all channels to assess co-occurance of lamin A/C and mCherry signals.

### Nuclear envelope protein assessment in micronuclei

Fixed cells with micronuclei were identified by scanning in DAPI. Relevant fields were recorded and imaged on a Nikon confocal microscope focused within a single, central plane through main nuclei. Quantification of nuclear envelope components was performed in ImageJ using the Radial Profile Plot plugin. Briefly, the plugin produces a profile plot of integrated fluorescent intensity measurements, for a series of concentric circles, as a function of distance from the central point. A standardized circular ROI was used to collect all micronuclear radial profile measurements, with the center of the ROI placed over the center of each micronucleus. Radial profile plots were collected in DAPI (to determine the distance cutoff for each micronucleus circumference) and in the lamin B1 channel (to quantify its abundance within the outer rim of the micronuclear envelope). Background subtracted lamin B1 measurements at the outer rim of each micronuclear envelope were recorded. This process was repeated to measure lamin B1 presence at the outer rim of primary nuclei and report the relative ratio of lamin B1 incorporation within nuclear envelopes of micronuclei and primary nuclei occurring in the same cell, for comparison. For primary nuclear measurements, ROI were arranged around the smallest circular ROI touching three sides of the main nucleus.

### Lagging chromosome measurements

Kif18A KO RPE-1 cells were synchronized with 10 μM CDK-1 inhibitor R0-3306 (Sigma-Aldrich) diluted in MEM-α media with 10% FBS and 1% antibiotic for 15 hours before drug was washed out. Cells were fixed 1 h post release to enrich for late anaphase cells. RPE-1 WT cells were treated with nocodazole (Sigma-Aldrich) at (5 μM) for 2 h before drug was washed out to induce merotelic attachments. Cells were fixed 45 min post washout to enrich for late anaphase cells.

Fixed cells in late anaphase were imaged in multiple z-stack focal planes to capture both focused spindle poles and individual ACA puncta. Image analysis was done in ImageJ with the straight line segment tool to obtain individual chromosome-to-pole distance measurements. One end of the line was anchored at the focused plane of a single spindle pole, and the other end was moved to the center of each focused ACA puncta (in the appropriate focal plane) for all 23 centromeres positioned for segregation to that pole (spindle-to-centromere distances taken for each respective half-spindle). All centromere positions were recorded, and average chromosome distribution was calculated for each individual half spindle measured, to account for cells fixed in different stages of anaphase. Late lagging chromosomes were determined as those centromeres with measured centromere to spindle distances greater than two standard deviations outside the averaged centromere to spindle distances for all centromere puncta per each half spindle. Normalized centromere to spindle distance were obtained for each late lagging chromosome by dividing that centromere’s centromere to spindle distance by the averaged centromere to spindle distance for that respective half spindle.

### Chromatin decondensation / Micronuclear envelope expansion measurements

RPE-1 cells were transfected with GFP-H2B plasmid. 24 h after transfection, cells were transferred to CO_2_–independent media for live imaging. For nocodazole or vehicle control DMSO conditions, RPE-1 cells were treated with drug or vehicle control for 2 h before drug washout (cells flushed 3x with PBS) immediately before transferring to CO_2_ –independent media and beginning filming. Cells found to be in metaphase were imaged every 2 min using a 40x objective and imaged for a duration of at least 4 h.

Chromatin decondensation measurements were obtained using ImageJ by applying the minimum threshold to completely cover chromatin (GFP-H2B) signal and measuring the area of the thresholded region corresponding to an individual chromatin mass forming either 1) a daughter nucleus or 2) a micronucleus. Frames which did not provide appropriate spatial separation to select only a single daughter cell nucleus or micronucleus without interference (overlapping other nuclei / edge of frame) were excluded from measurement. Area measurements began in the first frame of chromatin decondensation (following completion of chromatin segregation in late anaphase), and measurements were taken every subsequent frame for one-hour post anaphase onset, or until the chromatin area plateaued.

**Supplemental Figure 1:**
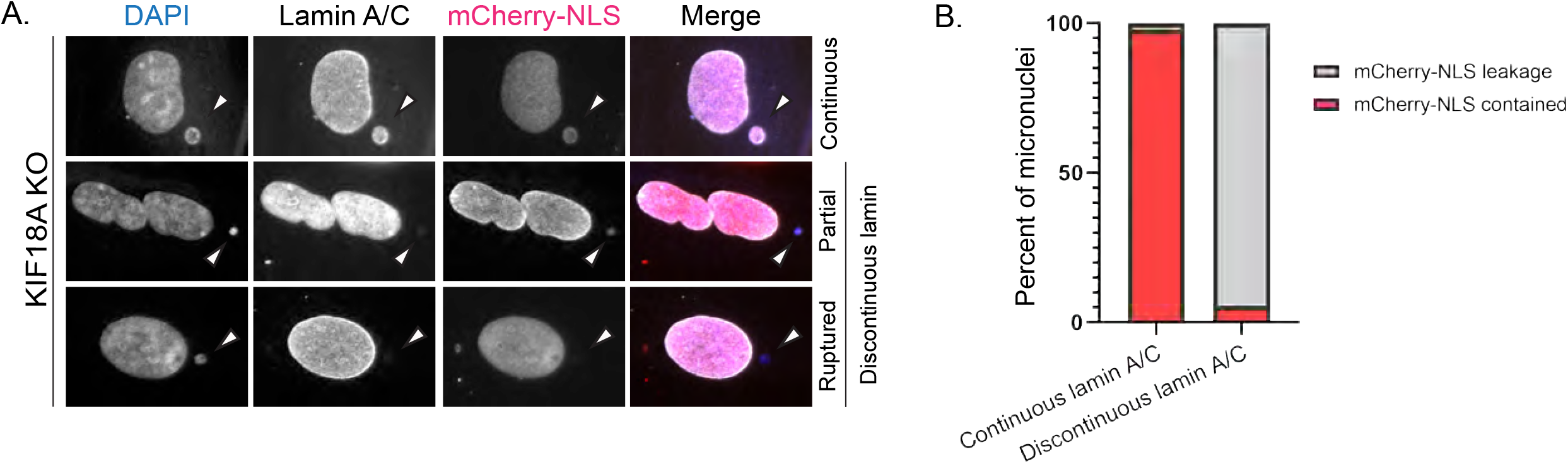
Validation of lamin A/C staining to assess micronuclear envelope integrity. **(A)** Representative images of KIF18A KO cells, and RPE1 cells treated with nocodazole washout, stained with DAPI (to identify micronuclei) and antibodies against lamin A/C and mCherry approximately 24 hours following transfection with mCherry-NLS construct. (B) Percentage of micronuclei exhibiting retainment or leakage of mCherry-NLS signal as a function of continuous or discontinuous nuclear membrane signal (lamin A/C). Of 140 micronuclei imaged and scored: 47 micronuclear envelopes showing continuous lamin A/C signal contained mCherry signal (1 showed mCherry leakage); 87 micronuclear envelopes showing discontinuous lamin A/C signal experienced mCherry leakage (5 retained mCherry signal). Comparison of these two criteria for the same micronuclei indicated that lamin A/C signal alone allowed us to correctly evaluate the integrity of the micronuclear envelope for 96% of cells (134/140 micronuclei). Validation data were collected from two independent experiments.

**Supplemental Figure 2:**
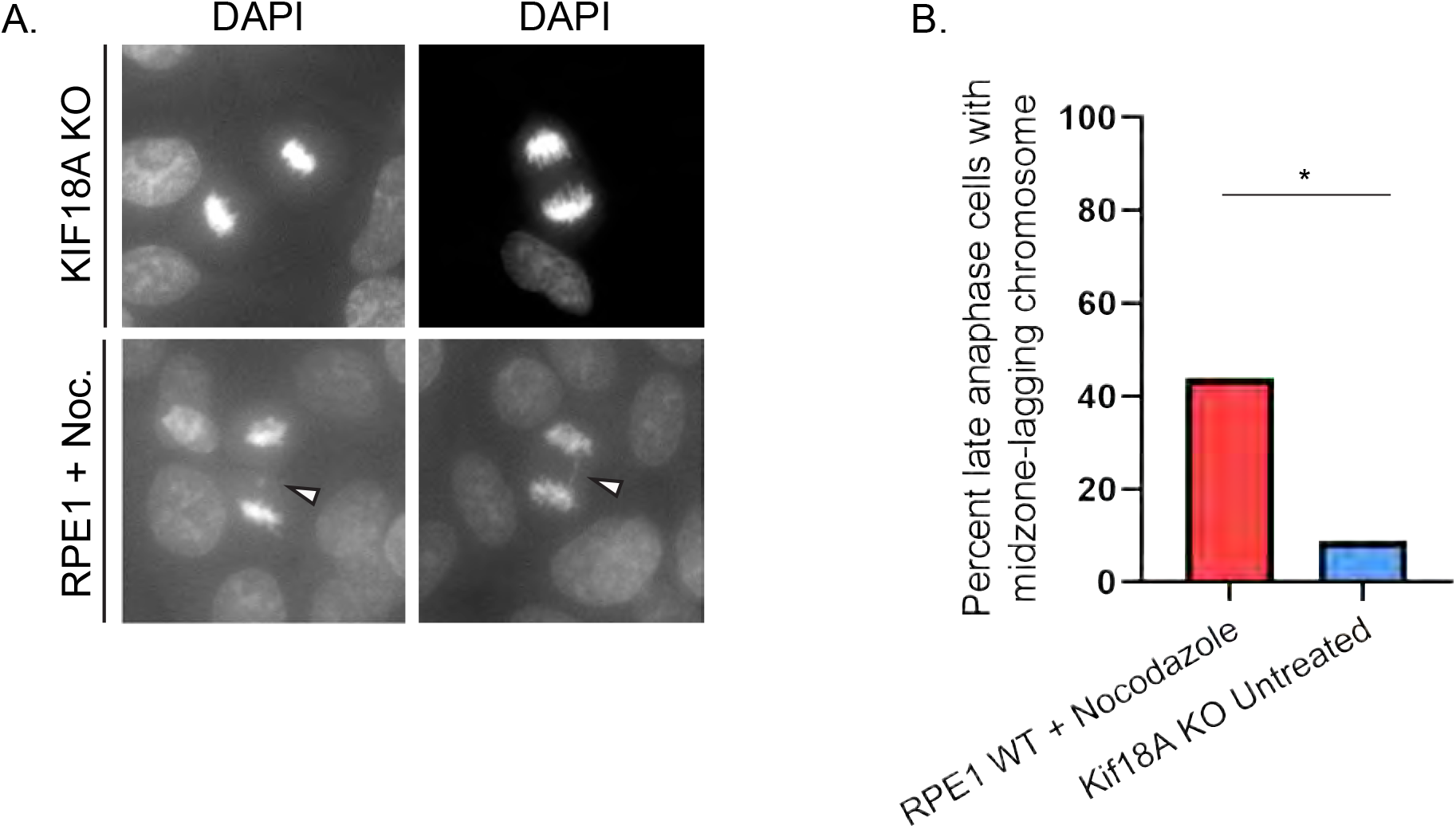
Late lagging chromosomes are infrequently observed in KIF18A KO late anaphase cells. **(A)** Representative late anaphase RPE1 cells treated with nocodazole washout or containing KIF18A KO mutations. Late-lagging chromosomes indicated via white arrowhead. **(B)** Percentage of late anaphase cells containing one or more lagging chromosomes from the indicated experimental conditions. Late-lagging chromosomes were observed in 44% (52/118) of nocodazole-washout treated RPE1 cells, and in 9% (4/43) of *KIF18A* KO cells; * p < 0.001. Data are from one experiment. Indicated p value was calculated by χ2 analysis.

**Supplemental Figure 3:**
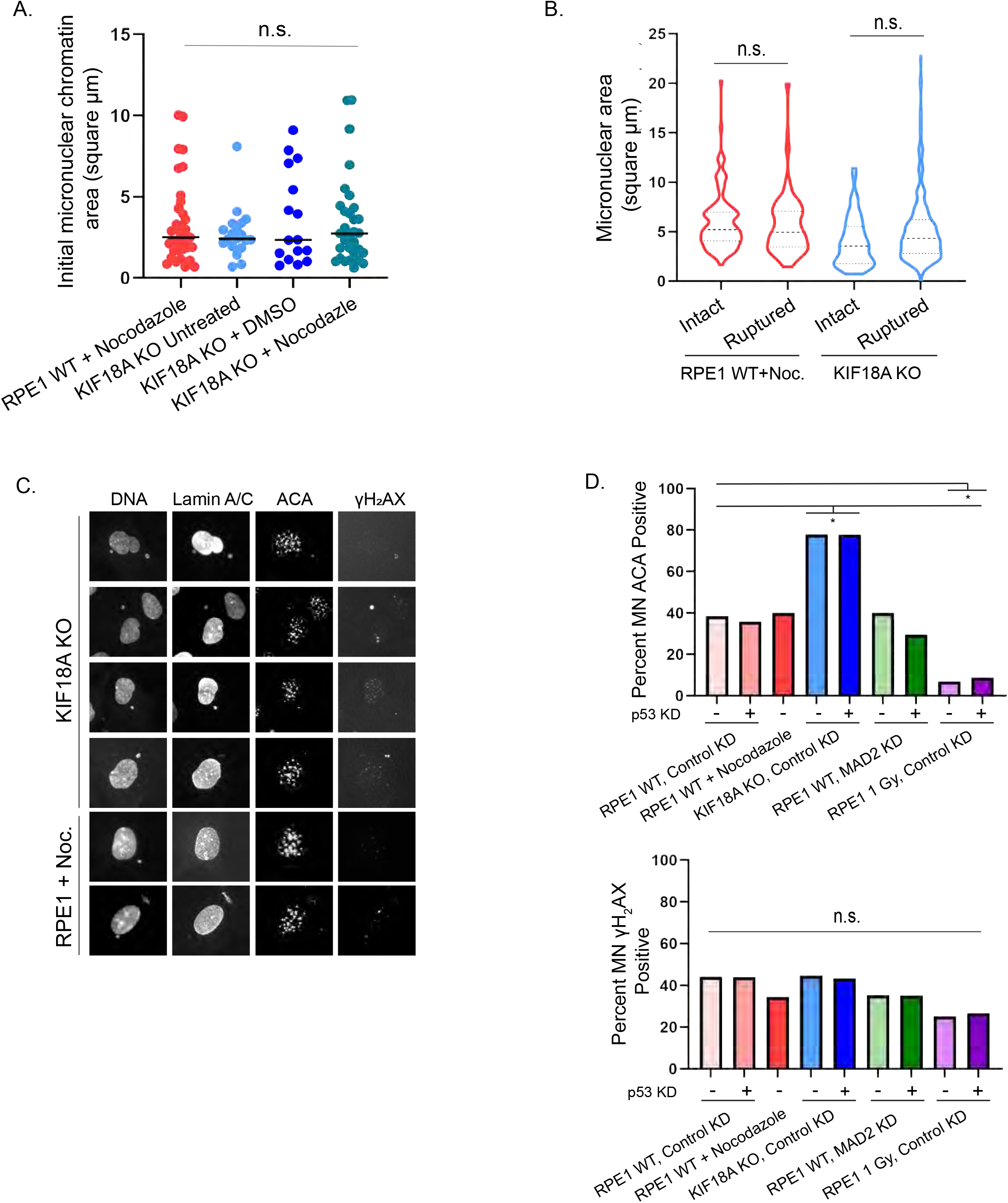
Micronuclear envelope rupture incidence does not strongly correlate with initial chromatin area, micronuclear area, centromere presence, or γH2AX DNA damage status. **(A)** Plot of initial chromatin area, a proxy for chromosome size, for micronuclei forming in KIF18A KO or RPE1 nocodazole-washout treated cells. n=35 (RPE1+nocodazole), n=19 (KIF18A KO untreated), n=31 (KIF18A KO+nocodazole), n=16 (KIF18A KO+DMSO), p = 0.7280, n.s. Data points indicate individual micronuclei. **(B)** Plot showing area of DAPI-stained micronuclear chromatin in an asynchronous, fixed-cell population, parsed by rupture-status (as determined via lamin A/C staining) for KIF18A KO or RPE1 nocodazole-washout treated cells. n= 219 (RPE1+nocodazole), n=310 (KIF18A KO). KIF18A KO intact vs. ruptured micronuclei, p = 0.063, n.s.; RPE1 + nocodazole intact vs. ruptured micronuclei, p = 0.73, n.s. **(C)** Representative images showing DAPI (DNA, to indicate micronuclei), along with antibodies against: lamin A/C (to assess micronuclear envelope integrity), anti-centromeric antibody (ACA; to assess centromere presence), and γH_2_AX (to assess DNA damage) associated with micronuclei arising in KIF18A KO cells and RPE1 nocodazole-washout treated cells. **(D)** Plot showing micronuclei positive for ACA signal (top graph; ACA signal indicates micronuclei likely contain whole chromosomes, while loss of centromeric signal suggests fragmentation), and γH_2_AX (bottom graph; γH_2_AX indicates foci of DNA damage), by method of micronuclear induction and p53 status. Data are pooled from three independent experiments. n=262 (RPE1 Control KD), n=304 (RPE1 Control+p53 KD), n=398 (KIF18A KO Control KD), n=359 (KIF18A KO Control KD+p53 KD) n=297 (RPE1 MAD2 KD), n=295 (MAD2 KD+p53 KD), n=312 (RPE1+nocodazole washout), n=518 (RPE1 1 Gy, Control KD), n=620 (RPE1 1 Gy, Control KD+p53 KD). *, p < 0.0001 (ACA), *, p < 0.0001 (γH_2_AX). Indicated p values for numerical data were obtained via unpaired Student’s t-test, for comparisons among two conditions, or a one-way ANOVA with post-hoc Tukey test, for comparisons among more than two conditions. Indicated p values for categorical data were calculated by χ^2^ analysis.

**Table. S1:** Counts of micronucleated cells (MN) *in vivo* as a fraction of total cells scored in healthy samples from thymus, spleen and liver tissues from the indicated genotypes. Reported values are the averages of two independent counts from these sampled tissues. Fixed tissues were labeled with Hoescht stain and lamin A/C antibodies.

|  | **Thymus** |  | **Spleen** |  | **Liver** |  |
| --- | --- | --- | --- | --- | --- | --- |
|  | MN / Total Cells | Ruptured / Total MN | MN / Total Cells | Ruptured / Total MN | MN / Total Cells | Ruptured / Total MN |
| *Kif18a<sup>gcd2/gcd2</sup>, Trp53<sup>+/+</sup>* | 83/1317<br>(6.3%) | 2/83<br>(2.4%) | 119/2587<br>(4.6%) | 6/74<br>(8.1%) | 57/1115<br>(5.1%) | 3/57<br>(5.3%) |
| *Kif18a<sup>gcd2/gcd2</sup>, Trp53<sup>+/tm1 Tyj</sup>* | 120/1496<br>(8.0%) | 14/120<br>(11.7%) | 68/1468<br>(4.6%) | 5/68<br>(7.4%) | 41/735<br>(5.6%) | 2/41<br>(4.9%) |
| *Kif18a<sup>gcd2/gcd2</sup>, Trp53<sup>tm1 Tyj/tm1 Tyj</sup>* | 36/602<br>(6.0%) | 4/36<br>(11.1%) | 97/2410<br>(4.0%) | 16/122<br>(13.1) | 46/811<br>(5.7%) | 8/46<br>(17.4%) |

**Table. S2:** Quantified micronuclear load within thymic lymphoma tissues from mice of the indicated genotypes. Fractions indicate number of micronucleated cells as a fraction of total cells scored, with percentage in parentheses. Individual micronuclear loads reported for each of three biological replicates per genotype. Far right column reports totals.

| Genotype | **Micronuclear load in thymic lymphoma** |  |
| --- | --- | --- |
|  | MN / Total Cells (%) | Total MN / Total Cells (%) |
| *Kif18a<sup>+/+</sup>, Trp53<sup>tm1 Tyj/tm1 Tyj</sup>* | 26/647<br>(4%) | 107/3099<br>(3.5%) |
|  | 37/1161<br>(3.2%) |  |
|  | 44/1291<br>(3.4%) |  |
| *Kif18a<sup>gcd2/gcd2</sup>, Trp53<sup>tm1 Tyj/tm1 Tyj</sup>* | 68/1286<br>(5.3%) | 253/4210<br>(6.0%) |
|  | 94/1272<br>(7.4%) |  |
|  | 91/1652<br>(5.5%) |  |

**Table. S3:** Counts of ruptured micronuclei within thymic lymphoma tumor tissues from the indicated genotypes, as determined via lamin A/C antibody staining. Frequency of ruptured micronuclei reported for each of three biological replicates per genotype. Fraction indicates ruptured micronuclei, of total micronuclei scored. Far right column reports totals.

| Genotype | Ruptured MN / Total MN (%) | Total Ruptured MN / Total MN (%) |
| --- | --- | --- |
| *Kif18a<sup>+/+</sup>, Trp53<sup>+/+</sup>* | 14/26<br>(54%) | 46/107<br>(43%) |
|  | 13/37<br>(35%) |  |
|  | 19/44<br>(43%) |  |
| *Kif18a<sup>gcd2/gcd2</sup>, Trp53<sup>tm1 Tyj/tm1 Tyj</sup>* | 29/68<br>(43%) | 117/253<br>(46%) |
|  | 47/94<br>(50%) |  |
|  | 41/91<br>(45%) |  |

**Table. S4:** Micronucleated RPE1 or KIF18A cells, as a fraction of total cells following each induction mechanism (see experimental design, Figure 4A). For both frequency of micronucleated cells and fraction of ruptured micronuclei described below in each condition, data are pooled from three independent experiments.

| MN Induction Method (RPE1) | MN / Total cells (%) |
| --- | --- |
| RPE1 Control KD | 43 / 4188 (1.0%) |
| RPE1 Control KD+p53 KD | 63 / 3536 (1.8%) |
| KIF18A KO Control KD | 35 / 661 (5.3%) |
| KIF18A KO Control KD+p53 KD | 48 / 869 (5.5%) |
| RPE1 MAD2 KD | 49 / 1223 (4.0%) |
| RPE1 MAD2 KD+p53 KD | 82 / 1157 (7.1%) |
| RPE1 + nocodazole washout | 223 / 4005 (5.6%) |
| RPE1 1 Gy, Control KD | 414 / 2189 (19.0%) |
| RPE1 1 Gy, Control KD+p53 KD | 700 / 3080 (23.0%) |

**Table. S5:**
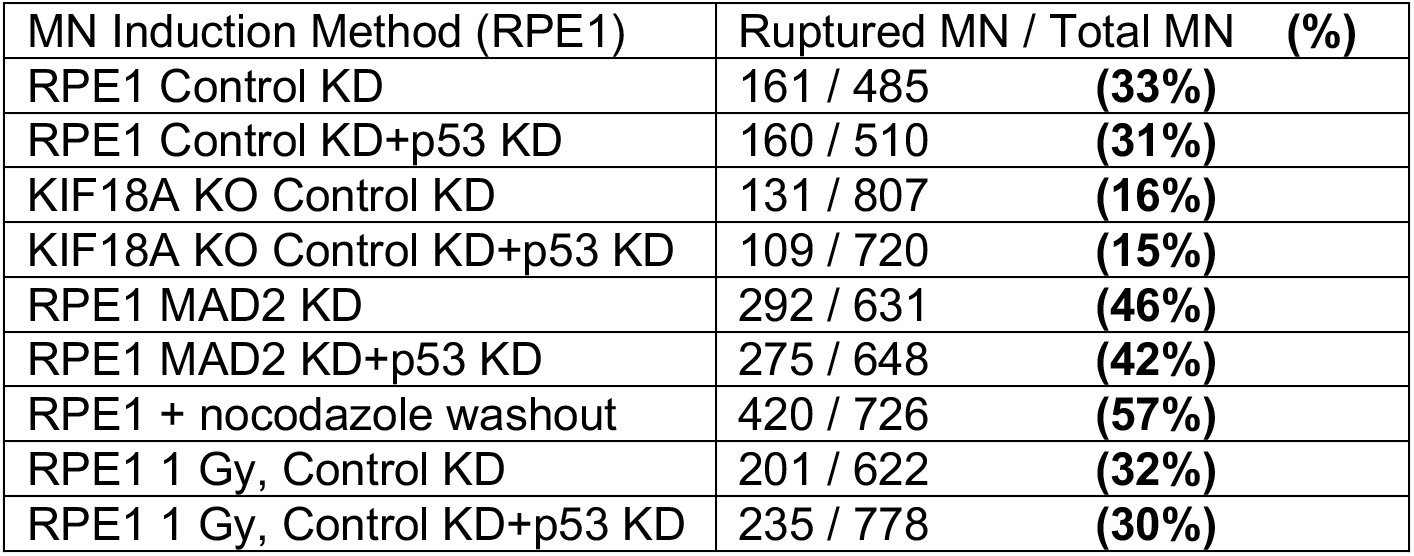
Frequency of micronuclear envelope rupture in RPE1 or KIF18A KO cells under the indicated conditions. Rupture assessed via lamin A/C staining. Data are pooled from four independent experiments.

| MN Induction Method (RPE1) | Ruptured MN / Total MN (%) |
| --- | --- |
| RPE1 Control KD | 161 / 485 (33%) |
| RPE1 Control KD+p53 KD | 160 / 510 (31%) |
| KIF18A KO Control KD | 131 / 807 (16%) |
| KIF18A KO Control KD+p53 KD | 109 / 720 (15%) |
| RPE1 MAD2 KD | 292 / 631 (46%) |
| RPE1 MAD2 KD+p53 KD | 275 / 648 (42%) |
| RPE1 + nocodazole washout | 420 / 726 (57%) |
| RPE1 1 Gy, Control KD | 201 / 622 (32%) |
| RPE1 1 Gy, Control KD+p53 KD | 235 / 778 (30%) |

**Table. S6:**
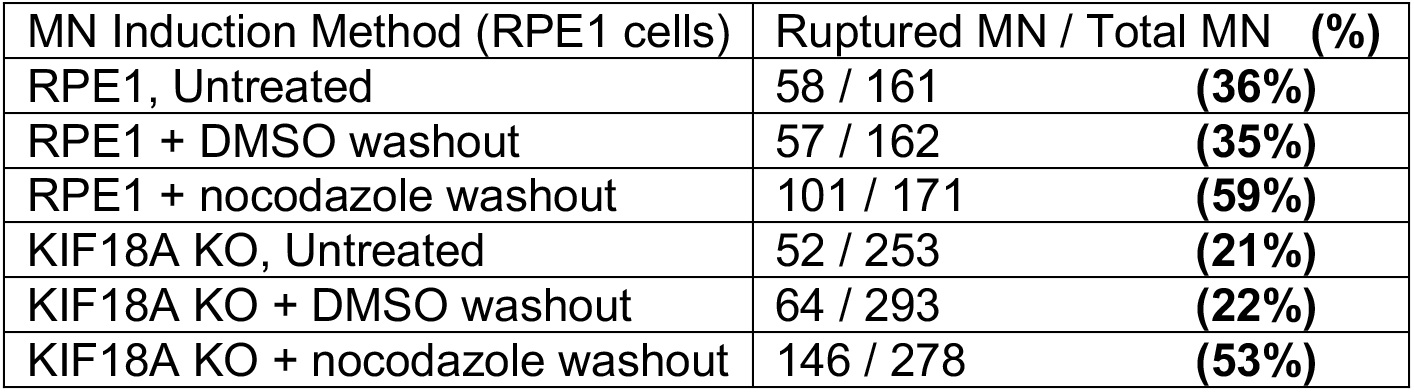
Micronucleated cells exhibiting micronuclear envelope rupture in RPE1 and KIF18A KO cells subjected to DMSO treatment or nocodazole-washout, as indicated. Data are pooled from three independent experiments.

**Table. S7:** Counts of micronuclei lacking lamin B as a fraction of total micronuclei counted in thymic lymphoma tumor tissues from the indicated genotypes. Data are reported for each biological replicate per genotype. Fractions indicate number of micronuclei lacking lamin B out of total micronuclei scored. Far right column reports totals from all 3 replicates.

| Genotype | Lamin B recruitment to micronuclei in thymic lymphoma |  |
| --- | --- | --- |
|  | MN without lamin B / Total MN (%) | Grand totals (%) |
| *Kif18a<sup>+/+</sup>*, *Trp53<sup>tm1 Tyj/tm1 Tyj</sup>* | 52/107 (49%) | 150/311 (48%) |
|  | 54/104 (52%) |  |
|  | 44/100 (44%) |  |
| *Kif18a<sup>gcd2/gcd2</sup>*, *Trp53<sup>tm1 Tyj/tm1 Tyj</sup>* | 72/159 (45%) | 200/387 (52%) |
|  | 43/77 (56%) |  |
|  | 85/151 (56%) |  |

## Acknowledgements

We thank Nicole Bouffard and the University of Vermont Microscopy Imaging Center for suggestions and support regarding the imaging of histological tissues. This work was supported by a Vermont Space Grant Consortium Fellowship awarded to LAS under NASA Cooperative Agreement NNX15AP86H. The Jackson Laboratory Cancer Center (P30CA034196) Pilot Award to LGR and JS, and NIH grant GM121491 and a University of Vermont Cancer Center pilot grant to JS. Confocal microscopy was supported by NIH award number 1S10OD025030-01 from the National Center for Research Resources.

